# Genomic determinants of protein abundance variation in colorectal cancer cells

**DOI:** 10.1101/092767

**Authors:** Theodoros I. Roumeliotis, Steven Paul Williams, Emanuel Gonçalves, Fatemeh Zamanzad Ghavidel, Nanne Aben, Magali Michaut, Michael Schubert, James C. Wright, Mi Yang, Clara Alsinet, Rodrigo Dienstmann, Justin Guinney, Pedro Beltrao, Alvis Brazma, Oliver Stegle, David J. Adams, Lodewyk Wessels, Julio Saez-Rodriguez, Ultan McDermott, Jyoti S. Choudhary

**Author notes:** **Co-first author**. **Co-senior author** (Lodewyk Wessels, Julio Saez-Rodriguez, Ultan McDermott, Jyoti S. Choudhary). **Corresponding author** (J.S.C.), (T.I.R.) (lead contact).

## Abstract

Assessing the extent to which genomic alterations compromise the integrity of the proteome is fundamental in identifying the mechanisms that shape cancer heterogeneity. We have used isobaric labelling and tribrid mass spectrometry to characterize the proteomic landscapes of 50 colorectal cancer cell lines and to decipher the relationships between genomic and proteomic variation. The robust quantification of 12,000 proteins and 27,000 phosphopeptides revealed how protein symbiosis translates to a co-variome which is subjected to a hierarchical order and exposes the collateral effects of somatic mutations on protein complexes. Targeted depletion of key chromatin modifiers confirmed the transmission of variation and the directionality as characteristics of protein interactions. Protein level variation was leveraged to build drug response predictive models towards a better understanding of pharmacoproteomic interactions in colorectal cancer. Overall, we provide a deep integrative view of the molecular structure underlying the variation of colorectal cancer cells.

**Highlights:**

- The cancer cell functional “co-variome” is a strong attribute of the proteome.
- Mutations can have a direct impact on protein levels of chromatin modifiers.
- Transmission of genomic variation is a characteristic of protein interactions.
- Pharmacoproteomic models are strong predictors of response to DNA damaging agents.

**Abbreviations:**

COREAD: Colorectal Adenocarcinoma
IMAC: Immobilized Metal ion Affinity Chromatography
ROC: Receiver Operating Characteristic
AUC: Area Under the Curve
WGCNA: Weighted Correlation Network Analysis
CNA: Copy Number Alteration
SOM: Self-Organizing Map
QTL: Quantitative Trait Loci
MSI: Microsatellite Instability
CPS: Colorectal Proteomic Subtypes

## Introduction

Tumours exhibit a high degree of molecular and cellular heterogeneity due to the impact of genomic aberrations on protein networks underlying physiological cellular activities. Modern mass spectrometry based proteomic technologies have now the capacity to perform highly reliable analytical measurements of proteins in large sizes of subjects and analytes providing a powerful tool in the quest for regulatory associations between genomic features, gene expression patterns, protein networks and phenotypic traits (Mertins et al., 2016; Zhang et al., 2014; Zhang et al., 2016).However, understanding how genomic variation leads to variable proteomic landscapes and distinct cellular phenotypes remains challenging due to the enormous diversity in the biological characteristics of proteins. Studying protein co-regulation holds the promise to overcome the challenges associated with molecular complexity and is now gaining ground in the study of molecular networks as it can efficiently predict gene functions (Stefely et al., 2016; Wang et al., 2016).Colorectal cancer cell lines are widely used as a model that approximates cancer behaviour in a variety of cellular and biochemical assays however their proteome based characteristics and the genomic factors underlying protein variation remain largely unexplored.

Here we leveraged the accurate quantification of a 12k total proteome obtained by the application of isobaric labelling and tribrid mass spectrometry analysis on a panel of 50 colorectal cancer cell lines, first to build *de novo* proteome-wide representations of biological functions inferred by protein co-variation, highly predictive of protein complexes and interactions, and second to rationalize the impact of genomic variation in the context of the cancer cell co-variation protein interaction network. We selected to study the colorectal cancer cell lines panel as it has been extensively characterised by whole exome sequencing, gene expression, copy number and methylation arrays, and the frequency of molecular alterations is similar to that seen in clinical cohorts (Iorio et al., 2016). The cancer cell functional “co-variome” appeared to be a strong attribute of the proteome, revealing the interdependencies of protein complexes and assigning putative functions to uncharacterized gene products. Additionally, our proteomics data were complemented by protein phosphorylation measurements encompassing a total of 27,000 phosphorylated sequences which demonstrated that co-variation of phosphorylation can also highlight known and novel biology. We assessed the direct effects of mutations on protein abundances and we integrated these effects with the cancer cell co-variome to uncover protein network vulnerabilities by the identification of possible collateral effects on protein complexes. Proteomic analysis of human iPS cells engineered with gene knockouts of key chromatin modifiers confirmed that genomic variation can be transferred from directly affected proteins to tightly co-regulated distant gene products through protein interactions. A significant number of drug response predictive models were uniquely attributed to protein level variation leading to a better understanding of pharmacoproteomic interactions in colorectal cancer. Our results constitute a comprehensive in-depth resource elucidating the molecular organization of colorectal cancer cells widely used in cancer research.

## Results

### Proteome and phosphoproteome coverage

To assess the extent of variation in protein and phosphorylation abundances within a panel of 50 colorectal cancer cell lines (COREAD) we utilized isobaric peptide labelling (TMT-10plex) and MS3 quantification (**Figure S1A**). We obtained relative quantification between the different cell lines in a log2 scale for 12,306 unique proteins and 27,423 non-redundant phosphopeptides at FDR<1% (9,489 and 12,061 in at least half of the samples respectively) (**Figure S1B**) (**Table S1**,**Table S2**). The average correlation between biological replicates was significantly higher than that of non-replicates (Pearson’s r=0.74 and −0.02 respectively, p-value=5.1e-69) which was also observed in the inter-laboratory comparison with previously published TMT data for six colorectal cancer cell lines (McAlister et al., 2014) (**Figure S1F**) confirming that subtle differences between the cell lines can be detected using our proteomics approach. Similar levels of global protein variation were observed across the cell lines with an average standard deviation of 1.7-fold denoting highly variable proteomes (**Figure S1C**). Correlation analysis between mRNA (publicly available data) and protein relative abundances across the cell lines indicated a significant, yet moderate concordance of the two molecular levels with an average Pearson correlation r=0.57 for matched cell line mRNA and protein data, and r=−0.015 for unrelated samples (Welch t-test, p-value < 2.2e-16) (**Figure S1G**). Overall, highly variable mRNAs tend to correspond to highly variable proteins (Spearman’s r=0.62) although with a wide distribution (**Figure S1H**). Notably, several genes including *TP53* displayed high variation at the protein level despite the low variation at the mRNA level, suggesting a significant contribution of post-transcriptional regulatory mechanisms to their total protein levels.

To identify genomic variants at the protein level we also searched our MS/MS spectra against a customized protein database containing all amino acid substitutions encoded by 77k missense mutations (Iorio et al., 2016). We identified 769 unique variant peptides mapped to 558 proteins (**Table S3**) (**Figure S1D**) including several mutated cell differentiation markers (CD44, CD46, ITGAV, ITGB1, ITGB4 and TFRC) that can be useful in targeted immunoaffinity based applications for discrimination of cancer cells as well as a range of mutated protein complexes (e.g spliceosomal, MCM, eIF3, CCT, Ksr1, Emerin). Two characteristic examples of mutant peptides for KRAS and CTNNB1 colorectal cancer genes are depicted in **Figure S1E**. Although the coverage in single amino acid variants is limited to proteins with medium to higher expression, it is concordant with the overall variant frequency and distinctly higher in hypermutated lines. Our proteogenomic search reveals the unique proteotypes of the COREAD cell lines.

### The subunits of protein complexes maintain tight stoichiometry of total abundance post-transcriptionally

As a means of simplifying the complexity of protein abundance variation we examined whether protein co-variation patterns detected across the cell lines could help aggregate the thousands of single protein measurements into a much smaller number of biologically meaningful clusters. The Spearman’s correlation coefficients between proteins with known physical interactions in protein complexes catalogued in the CORUM database (Ruepp et al., 2010) was bimodal-like and clearly shifted to positive values with mean 0.41. The respective distribution of all pairwise protein-to-protein comparisons displayed a normal distribution with mean 0.09 (**Figure S2A, left panel**). Specifically, 388 partially overlapping CORUM complexes, representing the most significantly correlating set, showed a greater than 0.4 median correlation between their components (**Table S4**). In contrast, the distribution of Spearman’s coefficients between CORUM pairs based on mRNA co-variation profiles was only marginally shifted towards higher correlations (**Figure S2A, right panel**). This indicates that the subunits of the complexes are tightly regulated post-transcriptionally. Indeed, comparative Receiver Operating Characteristic (ROC) curves showed that our proteomics data outperformed mRNA data in recapitulating protein complexes and STRING (Szklarczyk et al., 2015) interactions (CORUM ROC AUC: 0.77 vs 0.56, and STRING ROC AUC: 0.75 vs 0.58; for proteomics and gene expression respectively) (**Figure 1A and Figure 1B**). The ability to also recapitulate any type of STRING interaction indicates that protein co-regulation also encompasses functional relationships beyond structural physical interactions.

**Figure 1.**
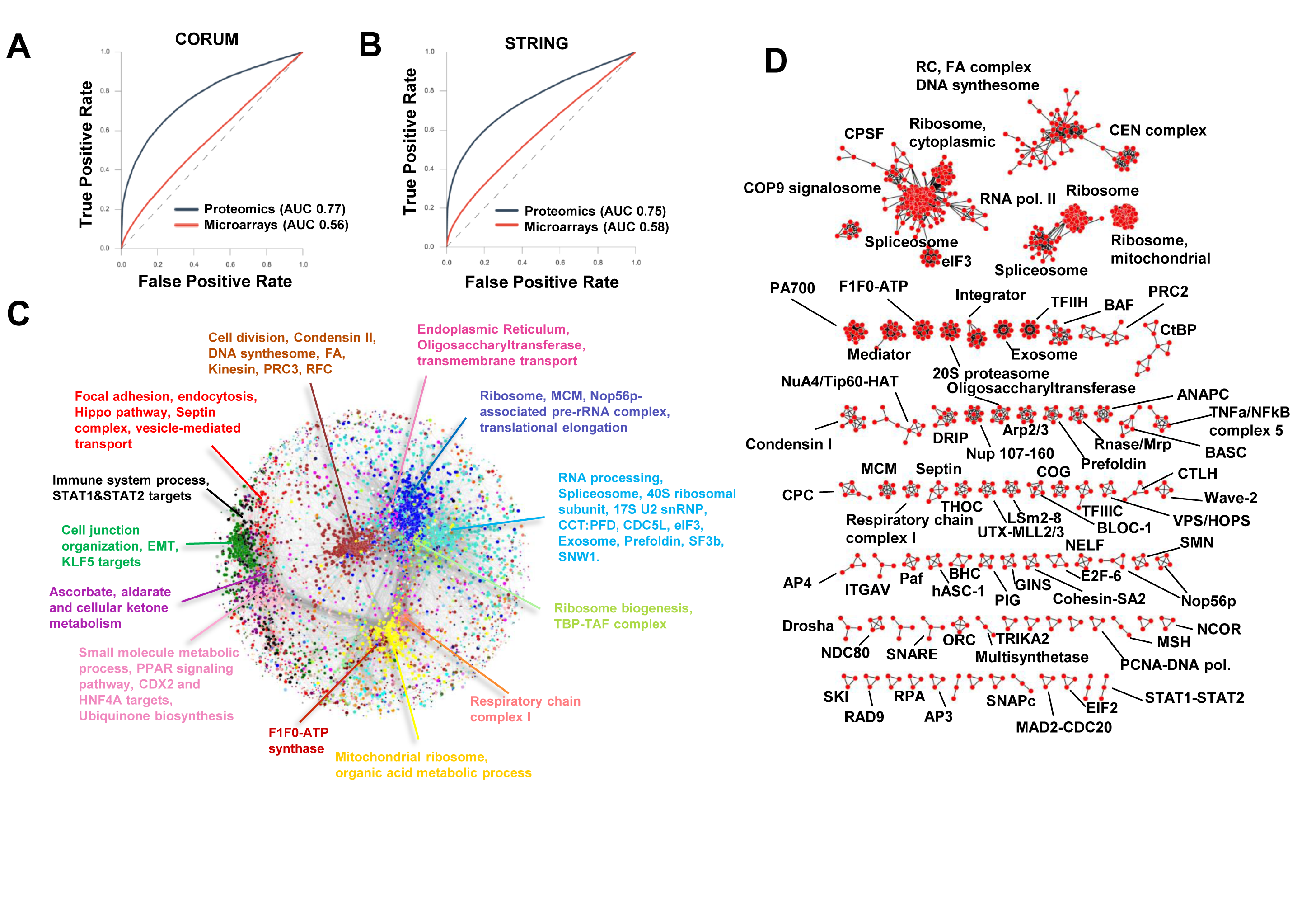
Protein co-variation networks in colorectal cancer cell lines. A) Receiver Operating Characteristic (ROC) curve illustrating the performance of proteomics and microarrays based correlations to predict known interactions from the CORUM protein complexes database and from B) the STRING database. C) The basic structure of the full WGCNA network. Protein modules are color-coded and representative enriched terms are used for the annotation of the network. D) Protein abundance correlation networks derived from WGCNA analysis for enriched CORUM complexes.

### Weighted correlation networks reveal the interdependencies of protein complexes and biological processes

We have shown above that the co-regulation of protein abundance is a strong predictor of physical and functional associations. We therefore conducted systematic genome-wide analysis of the colorectal cancer cell protein-protein correlation network. To this end, we performed a weighted correlation network analysis (WGCNA) (Langfelder and Horvath, 2008) using 8,469 proteins quantified in at least 80% of the cell lines. A total of 203 modules of co-regulated proteins ranging in size from 3 to 1,065 proteins (median = 9) were detected. A comprehensive description of the modules was devised based on enrichment analysis (**Table S5**) and the basic structure of the colorectal cancer network is depicted in **Figure 1C**. We found that approximately 60% of the modules displayed overrepresentation of protein complexes (**Figure 1D**) and that the largest modules were associated with RNA processing, plasma membrane, cytosolic ribosome, cell cycle, mitochondrial translation, mitochondrial respiratory chain, immune response and small molecule metabolic process (FDR<0.01). The full WGCNA network with weights greater than 0.02 is provided in **Table S6**.

To identify regulators of protein modules not explained by physical protein interactions we examined whether enriched transcription factors from ENCODE and ChEA databases in a given module were indeed co-expressed at the protein level along with their transcriptional targets. We found that the “small molecule metabolic process” module was enriched for the transcription factors HNF4A and CDX2 with 66 and 22 transcriptional targets respectively (Benj. Hoch. FDR=0.00027 and FDR=0.00012 respectively). HNF4A (Hepatocyte nuclear factor 4-alpha) is an important regulator of metabolism, cell junctions and the differentiation of intestinal epithelial cells (Garrison et al., 2006) and has been previously associated with colorectal cancer proteomic subtypes in human tumours analyzed by the CPTAC consortium (Zhang et al., 2014). The “plasma membrane” module included 60 transcriptional targets of KLF5 (Benj. Hoch. FDR=0.00174) which itself was a member of this module. KLF5 is predominantly expressed in the proliferating cells of the crypt and appears to play a growth regulatory role in the intestine (Bateman et al., 2004). Interestingly, KLF5 was significantly correlated with HNF4A (Pearson=0.79, Benj. Hoch. FDR=4.17E-10) providing a potential link between cell communication and HNF4A regulated metabolic functions. Moreover, the “plasma membrane” module was enriched for an epithelial-mesenchymal transition (EMT) gene set and was characterized by the anti-correlation between the epithelial marker CDH1 and the mesenchymal marker Vimentin (VIM). STAT1 and STAT2 are master regulators of the “immune response” module with a total of 45 targets including several interferon-induced proteins (Benj. Hoch. FDR=5.06E-17 and FDR=2.05E-21 respectively). The protein correlation modules clearly serve as unique attributes by which upstream regulatory events can be identified at the protein level. This approach leverages a greater number of proteomic features and extends our knowledge about cancer associated regulators beyond the use of profiles for single transcription factors.

To better understand the interdependencies of protein complexes and biological processes of the colorectal cancer cells in a global way we plotted the module-to-module relationships as a correlation network. The nodes denote significant terms from each module and the edges represent pairwise correlations between the eigengenes (first principal component, **Table S7**) of the modules (Pearson>0.46, Benj. Hoch. FDR<0.01) (**Figure S2B**). The topology of this network revealed a strong coordination between the cytoplasmic ribosome and the RNA processing complexes that were further linked with protein processing complexes as well as with cell division and DNA repair protein complexes. Interestingly, the MCM complex, in contrast to the GINS complex, was better correlated with the cytoplasmic ribosome rather than the cell division processes coinciding with the proposed mechanisms which couple DNA and protein syntheses (Berthon et al., 2009). The 26S proteasome complex was directly associated with the RNA spliceosome generating the hypothesis of another un-expected interplay between RNA processing and protein turnover. In fact, the mechanistic connections between transcription and the Ubiquitin-Proteasome system have been previously discussed (Muratani and Tansey, 2003). The mitochondrial functions formed a distinct cluster comprised of mitochondrial translation and cellular respiration complexes. The HNF4A-metabolic related module is linked with the mitochondrial cluster as many of the HNF4A targets are involved in mitochondrial processes. Protein processing and trafficking complexes were grouped together and were associated with actin related complexes as well as with proteins involved in antigen presentation. The immune response signature was tightly correlated with focal adhesion, caveola proteins, septin complex and key proteins of the Hippo signalling pathway such as SAV1, STK3 and YAP1 revealing novel associations. Highlighting the modules with high mean mRNA-to-protein correlations on the network confirmed that the HNF4A, CDX2, KLF5, STAT1 and STAT2 modules were strongly driven by transcription, whereas nodes representing protein complexes corresponded poorly to mRNA levels (**Figure S2B**). The correlation of RNA to protein levels also appears to be modestly influenced by protein class (**Figure S3A**). Interestingly, proteins characterized by degradation that is best explained by a two-state model with two different degradation rates (non-exponentially degraded, NED) present significantly lower mRNA-to-protein correlations compared to their exponentially degraded (ED) counterparts (McShane et al., 2016) that are explained by one-state model (**Figure S3B**).

Examination of unannotated modules revealed correlation between tRNA methyltransferases and eukaryotic translation elongation factors (**Figure S3C**). This suggests that the maintenance of total stoichiometry is also a feature of interactions between protein complexes. We also mapped 66 uncharacterized proteins to 30 modules that allowed them to be functionally annotated based on their associated module function. Some candidates for further investigation include: cell cycle (C1orf112, C14orf80, C12orf45, C9orf78, C10orf12, C1orf52), tRNA processing (C18orf21), mitochondrial translation (C6orf203, C2orf47), chaperonin-containing T-complex (C12orf29), N-terminal protein amino acid acetylation (C8orf59), protein N-linked glycosylation via asparagine (C20orf24), Oxidative phosphorylation (C8orf82), ribonuclease H2 complex (C8orf76) and BLOC-1 complex (C10orf32). Taken together, our protein correlations reveal a higher order of cellular functions in a well-organized structure shaped by the compartmental interactions between protein complexes and clearly divided into transcriptionally and post-transcriptionally regulated sectors.

### *De novo* prediction of phosphorylation networks reveals novel functional relationships

The scale of global phosphorylation survey accomplished here offers the opportunity for the *de novo* prediction of kinase-substrate associations inferred by co-changing phosphorylation patterns that involve kinases (Ochoa et al., 2016; Petsalaki et al., 2015). Phosphorylated proteins are highly enriched for spliceosomal and cell cycle functions and cover a range of cancer related pathways (**Figure S3D**). Notably, for about 450 partially overlapping CORUM complexes more than 60% of their subunits were found to be phosphorylated. To detect differential phosphorylation we regressed protein abundances from the respective phosphorylation profiles as the two levels of information are strongly correlated (**Figure S3E**). Pairwise correlation analysis among 213 variable phosphopeptides belonging to 144 kinases, and the 787 most variable phosphopeptides from other protein types (**Table S8**) revealed a strong enrichment of nucleosome assembly proteins (mainly histones) (FDR=5.09E-08) correlating with three kinases, namely CSNK1A1, CDK13 and VRK3 (**Figure S4A**). CSNK1A1 is a casein kinase that participates in Wnt signalling where it is essential for β-Catenin phosphorylation and degradation (Liu et al., 2002). CSNK1A1 has also been implicated in the segregation of chromosomes during mitosis and may be cell cycle-regulated (Brockman et al., 1992). VRK3 has recently been shown to be an active kinase as well as a signalling scaffold in cells, with a specific role in DNA replication and chromatin dissociation during interphase (Park et al., 2015). Interestingly, VRK3 phosphorylation status was strongly correlated with the phosphorylation of 7 histones of which H2AFX presented the strongest correlation in spite of poor correlation of their respective protein abundance profiles (**Figure S4B, top panel**). Strongly maintained co-phosphorylation was also observed between RAF1, MAPK1, MAPK3 and RPS6KA3 of the MAPK pathway (**Figure S4C, left panel**) as well as between CDK1 and CDK7 of the cell cycle pathway (**Figure S4C, right panel**). The correlation plots of MAPK1 and MAPK3 phosphorylation and total protein are depicted in **Figure S4B, bottom panel**. Overall, these examples demonstrate that functional relationships are encrypted in the patterns of co-regulated phosphorylation events.

### Protein abundance and phosphorylation variation are associated with genomic alterations

Assessing the impact of non-synonymous protein coding variants and copy number alterations on protein abundance is fundamental in understanding the link between cancer genotypes and the dysregulated biological processes. To investigate this, we first examined whether driver mutations in any of the 17 colorectal cancer driver genes (Iorio et al., 2016) with at least 5 occurrences across the cell lines could alter the levels of their protein products. Strikingly, for 7 such genes (*PTEN, B2M, CD58, PIK3R1, ARID1A, BMPR2* and *MSH6*) we found that driver mutations had a significant negative impact on the respective protein abundances, in line with their function as tumour suppressors, whereas missense mutations in *TP53* were associated with elevated protein levels (ANOVA test, permutation-based FDR<0.05) (**Figure 2A**). Protein abundance variation of APC could not be systematically attributed to genomic variants although it is possible that extreme protein changes in individual cell lines could be the result of specific mutations. For example, the low relative abundance of APC protein in 5 out of 7 cell lines with log_2_Ratio<-0.8 could be explained by the presence of frameshift and nonsense mutations in these particular cell lines. Distinctly, for the majority of driver mutations in oncogenes, there was no clear relationship between the presence of mutations and protein expression. From these observations we conclude, that mutations in canonical tumour suppressor genes predicted to cause premature stop codons and ultimately nonsense-mediated decay of transcript were significantly associated with decreased protein abundance compared to driver mutations in oncogenes (Log_2_ mean protein abundance −0.65 vs +0.45 respectively) (**Figure 2B**), suggesting that these have distinct regulation or gain new function upon mutation.

**Figure 2.**
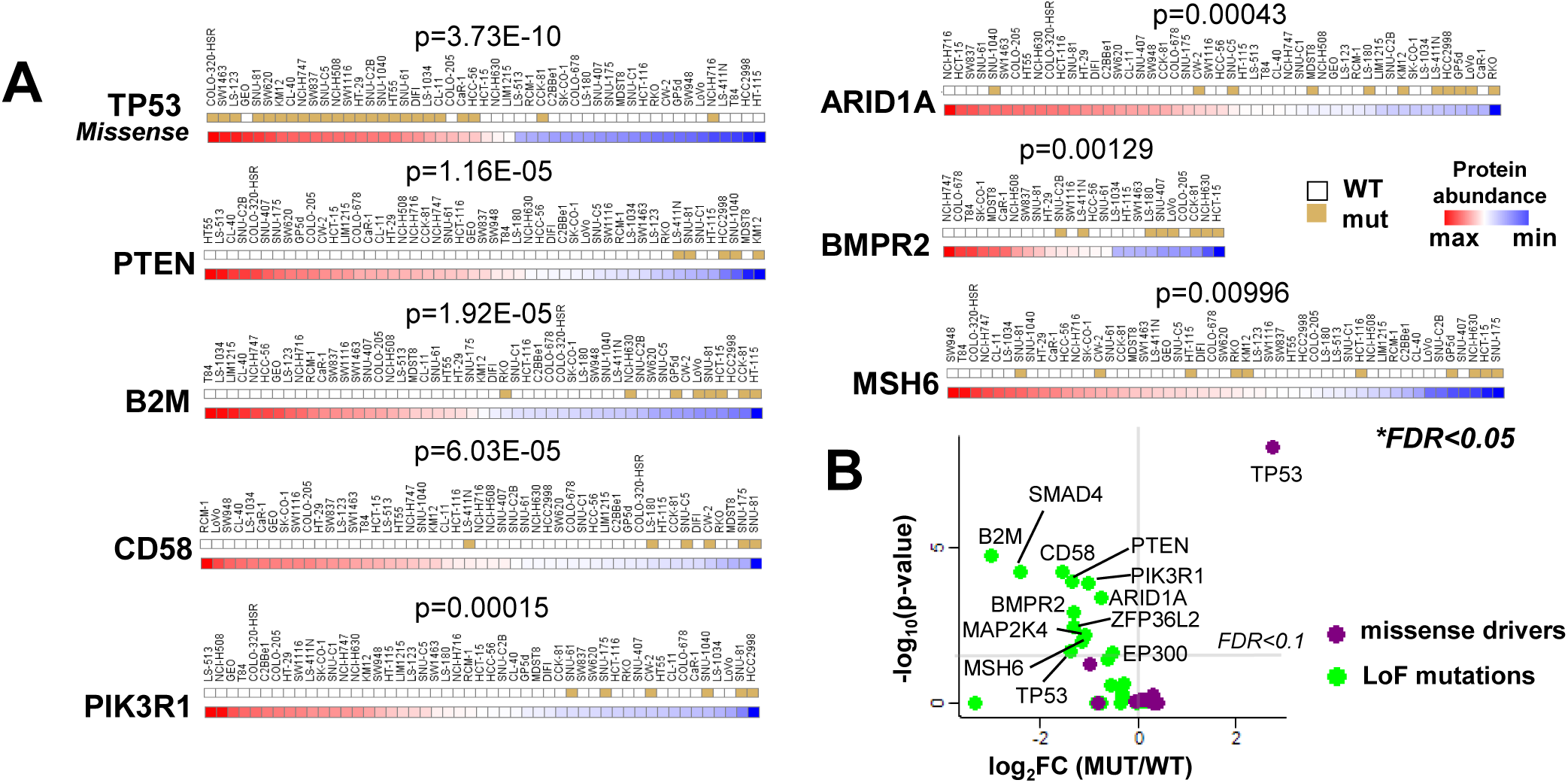
The effect of colorectal cancer driver mutations on protein abundances. A) Association of driver mutations in colorectal cancer genes with the respective protein abundance levels (ANOVA test, permutation based FDR<5%). The cell lines are ranked by protein abundance and the bar on the top indicates the presence of mutations with brown mark. B) Volcano plot summarizing the effect of loss of function (LoF) and missense driver mutations on the respective protein abundances.

We extended our analysis to globally assess the effect of mutations on the protein abundances. For 5,498 genes harbouring any type of non-synonymous protein coding variants in at least three cell lines, 626 proteins exhibited lower (N=566) or higher (N=60) abundances in the mutated versus the wild-type cell lines at ANOVA p-value<0.05 (all 77 proteins with FDR < 0.1 were associated with decreased levels of expression) (**Figure 3A**). This high confidence subset was enriched for phosphatidylinositol signalling proteins (KEGG FDR=0.0307: PTEN, PIK3R1, PIK3C2A, MTM1 and PIK3C2B) and included 5 tumour suppressors (MLH1, MSH2,NF1, PIK3R1, PTEN). Restricting the analysis to frameshift mutations only (the second most frequent mutation type), we found that 136 of the 389 genes presented lower abundances in the mutated cell lines with ANOVA p-value<0.05 of which 121 passed the 10% FDR cut-off (**Figure 3B**). Notably, the significantly affected proteins were strongly enriched for chromatin modification proteins (FDR=2.66E-10, N=23) and included 10 oncogenes and 4 tumour suppressors. The STRING network of the most significant hits is depicted in **Figure 3E**. A less pronounced impact of frameshift mutations was found at the mRNA level where only 15% of the 349 genes (with both mRNA and protein data) exhibited altered mRNAs abundances in the mutated samples at ANOVA p-value<0.05, only 19 of these were below the 10% FDR (**Figure 3C**). The overlap between the different analyses is depicted in **Figure 3D**. Considering all proteins negatively affected by mutations we found overrepresentation of proteins with certain domains (e.g. helicase) as well as enrichment of certain classes of enzymes such as kinases, transferases and hydrolases (**Figure 3F**) highlighting the protein classes that are subject to protein abundance reduction upon structural changes. Notably, 59 out of the 677 genes affected by genomic variants at p-value<0.05 are currently catalogued in the COSMIC Census list of genes for which mutations have been causally implicated in cancer.

**Figure 3.**
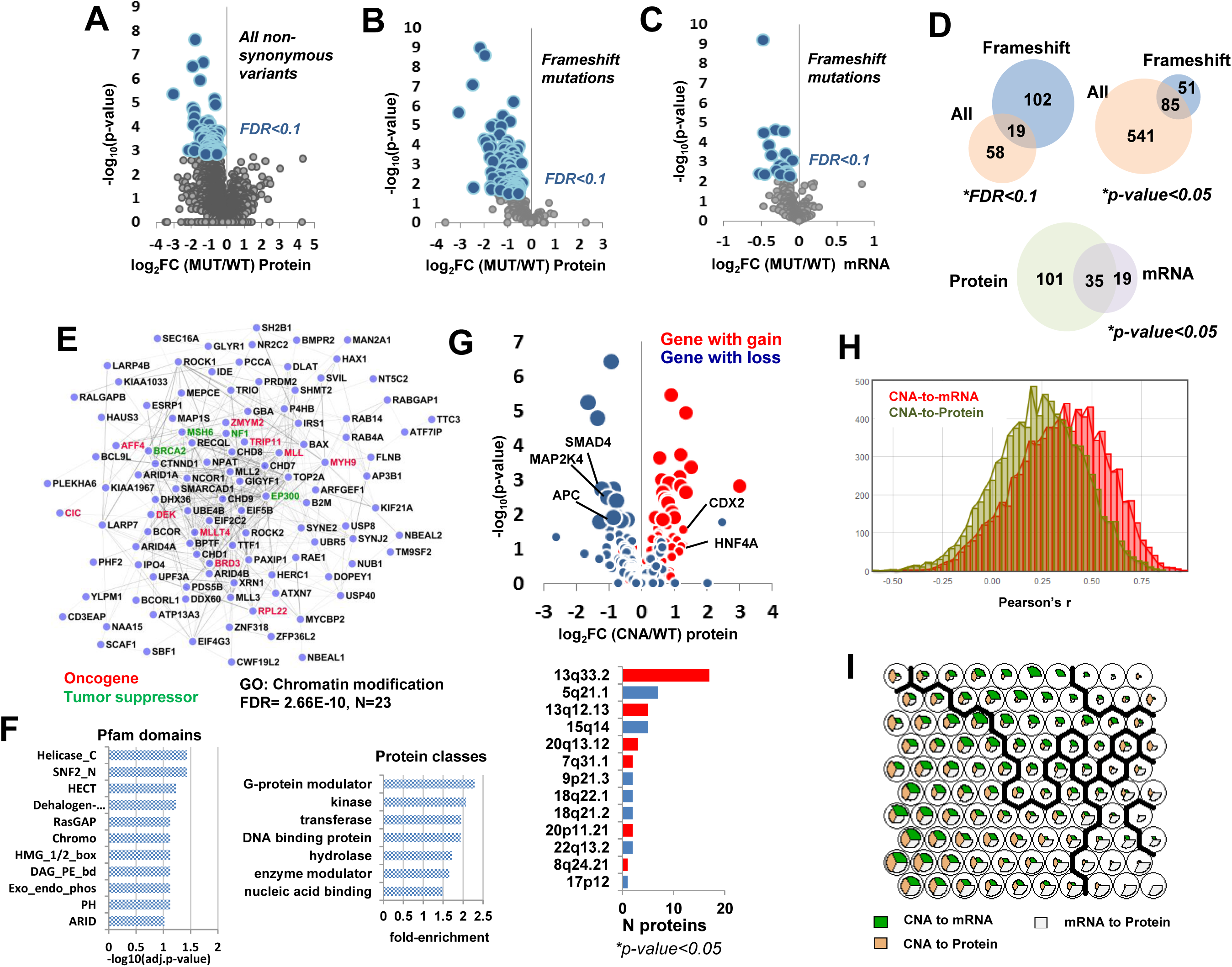
The global effects of genomic alterations on protein and mRNA abundances. A) Volcano plot summarizing the effect of all protein coding non-synonymous variants on the respective protein abundances (ANOVA test). B) Volcano plot summarizing the effect of all protein coding frameshift mutations on the respective protein abundances (ANOVA test). C) Volcano plot summarizing the effect of all protein coding frameshift mutations with both mRNA and protein measurements on the respective mRNA abundances (ANOVA test). D) Venn diagrams displaying the overlap between proteins affected by all types of mutations and proteins affected by frameshift only mutations at different confidence levels (top panel) and the overlap between proteins and mRNAs affected by frameshift mutations at p-value<0.05 (bottom panel). E) STRING network of the proteins down-regulated by frameshift mutations (permutation based FDR<0.1). F) Overrepresentation of Pfam domains (left panel) and PANTHER protein classes (right panel) for proteins negatively affected by mutations. G) Volcano plot summarizing the effect of recurrent copy number alterations on the protein abundances of the contained genes (binary data, ANOVA test). Red and blue points highlight genes with amplifications and losses respectively. Enlarged points highlight genes at FDR<10%. The bar plot (bottom panel) illustrates the number of affected proteins per genomic locus. Red and blue bars indicate amplifications and losses respectively. H) Distributions of the Pearson’s correlation coefficients for CNA to mRNA (red) and CNA to protein (green) correlations considering all genes across 38 cell lines with normalized log2 copy number values (source: Cancer Cell Line Encyclopedia). I) Self-Organizing Map trained on the Pearson correlation between CNA, mRNA and protein levels per gene across the cell lines. The fan plot within each neuron displays the magnitude of each one of the three vectors. Three main regulatory routes can be distinguished: good concordance between the three levels (left side cluster), CNAs corresponding to mRNAs but buffered at the protein level (top-middle cluster) and proteins well corresponding to mRNA irrespective of the presence of CNAs (bottom right cluster).

We also explored the effect of 20 recurrent copy number alterations (CNAs) using binary-type data on the protein abundances of 212 falling within these intervals. Amplified genes tended to display increased protein levels whereas gene losses had an overall negative impact on protein abundances although with several exceptions(**Figure 3G, top panel**). The 51 significant genes with ANOVA p-value <0.05 (31 genes at FDR<0.1) were mapped to 13 genomic loci. The 13q33.2 amplification encompasses the highest number of affected proteins (**Figure 3G, bar plot**). Losses in 18q21.2, 5q21.1 and 17p12 loci are associated with reduced protein levels of three important colorectal cancer drivers, SMAD4, APC and MAP2K4 respectively (FDR<0.1). Increased levels of CDX2 and HNF4A were modestly associated with 13q12.13 and 20q13.12 amplifications (p-value<0.1, FDR<30%). Global correlation using normalized log2 copy number ratios obtained from the Cancer Cell Line Encyclopaedia showed median CNA to mRNA and protein correlations 0.35 and 0.23 respectively (**Figure 3H**). To summarize the possible levels of regulation we trained a Self-Organizing Map (SOM) using the Pearson correlations coefficients between CNA and mRNA, CNA and protein, mRNA and protein (three vectors) for each protein, which indicated three main regulatory routes: good concordance between the three levels (**Figure 3I, left side cluster**), CNAs corresponding to mRNAs but buffered at the protein level (**Figure 3I, top-middle cluster**) and proteins well corresponding to mRNA irrespective of CNAs (**Figure 3I, bottom right cluster**). Taken together, our results show that copy number alterations more often affect the mRNA levels than the protein levels, which needs to be taken under consideration when gene expression data are used as a proxy of the protein levels for the identification of actionable pathways. A summary of all proteins affected by mutations and recurrent CNAs is in **Table S9**.

Next we assessed the direct impact of mutations on net protein phosphorylation. We found 72 differentially phosphorylated proteins in the mutated cell lines (Welch’s t-test, FDR<10%) with both positive and negative effects (**Figure S5A**). The SRC kinase and the RUNX1 transcription factor were among the top over-phosphorylated proteins while APC was among the top hypo-phosphorylated proteins. We then focused on eight colorectal cancer genes (*APC, TP53, KRAS, BRAF, PIK3CA, PTEN, RNF43* and *PIK3R1*) to individually assess the extended effects of driver mutations on the phosphorylation status of the colorectal cancer pathway. We found that *APC* mutations were associated with decreased phosphorylation of APC and increased phosphorylation of AXIN1, and that *PTEN* mutations were related to increased TP53 phosphorylation at 10% FDR (ANOVA test) (**Figure S5B**). Mutations in *PIK3CA* were associated with increased inactivating phosphorylation of BAD, while *BRAF* V600E mutants exhibited increased AKT1 and decreased ARAF phosphorylation (**Figure S5B**). As expected, for all the associations the respective total protein levels were undifferentiated (**Figure S5C**). These observations indicate a sophisticated level of cross talk between cancer genes.

Overall, we show that not all driver mutations have the same effect on protein abundance. We identify key mutations that significantly impact abundance levels of proteins, which converge in certain protein classes. We conclude that for only a small portion of the proteome the variation in abundance can be directly explained by mutations and that driver mutations also alter the phosphorylation status of colorectal cancer proteins.

### The consequences of genomic alterations extend to protein complexes

As tightly controlled maintenance of protein abundance appears to be pivotal for protein complexes and interactions, we hypothesize that genomic variation can be transferred from directly affected genes to distant gene protein products through protein interactions there by explaining another layer of protein variation. We retrieved strongly co-regulated interactors of the affected proteins and constructed mutation-vulnerable protein networks, comprised of 1,108 total protein nodes (**Figure 4A**) encompassing at least 25 protein complexes. One characteristic example was the BAF complex characterized by disruption of ARID1A protein abundance. Driver mutations in ARID1A were also significantly associated with decreased levels of the respective module (p-value = 0.01707) (**Figure 4B**) indicating a central role of ARID1A in the regulation of the profile of the complex. The WGCNA networks also revealed a correlation between the BAF complex and the functionally related PBAF complex containing the ARID2 and PBRM1 proteins which were however mapped to different modules (**Figure 4C**). We noticed that the PBRM1 sub-network displayed unusually poor overlap with STRING interactions, and the correlations were attributed to the effects of co-occurring mutations on the protein abundances, specifically in the hypermutated HT-115 cell line (**Figure S6A**). This indicates that in addition to functional relationships, protein co-regulation can also classify the effect of co-occurring genomic variants. These events are infrequent and observed on small modules, which lack functional connection between their components. To confirm whether the down-regulation of ARID1A, ARID2 and PBRM1 can indeed affect the abundance levels of their interactors we performed proteomic analysis on the respective CRISPR-Cas9 knockout (KO) clones derived from human iPS cells (**Table S10**). Down-regulation of ARID1A coincided with diminished levels of 8 partners in the predicted interactome that were closer to the core of the network and were known components of the BAF complex whilst more distant interactors were not affected (**Figure 4D**). This provides an indication that the topology of the correlation network can predict the relative strengths of interactions. Reduced levels of ARID2 resulted in the down-regulation of all three direct interactors (BRD7, PHF10 and SCAF11) and the significant loss of PBRM1 protein. Four components of the BAF complex were also weakly compromised in the ARID2 KO reflecting the overlap between the BAF and PBAF complexes. On the other hand, loss of PBRM1 had no effect on ARID2 or any of its interactors demonstrating that collateral effects transmitted through protein interactions can be directorial. Distinctly, loss of PBRM1 had no impact on the abundance of the constituents of its module confirming that the co-variation here is due to co-occurring genomic variation rather than direct interactions. Pathway enrichment analysis on the changing proteins detected in the KO cell lines, revealed the differential regulation of a number of biological processes reflecting the modulation of a wide range of target genes (**Figure 4E**). Notably, down-regulation of ARID2 specifically activated the MAPK pathway, actin cytoskeleton, ubiquitin proteolysis and immune related signalling pathways that were not affected by ARID1A and PBRM1 depletion.

**Figure 4.**
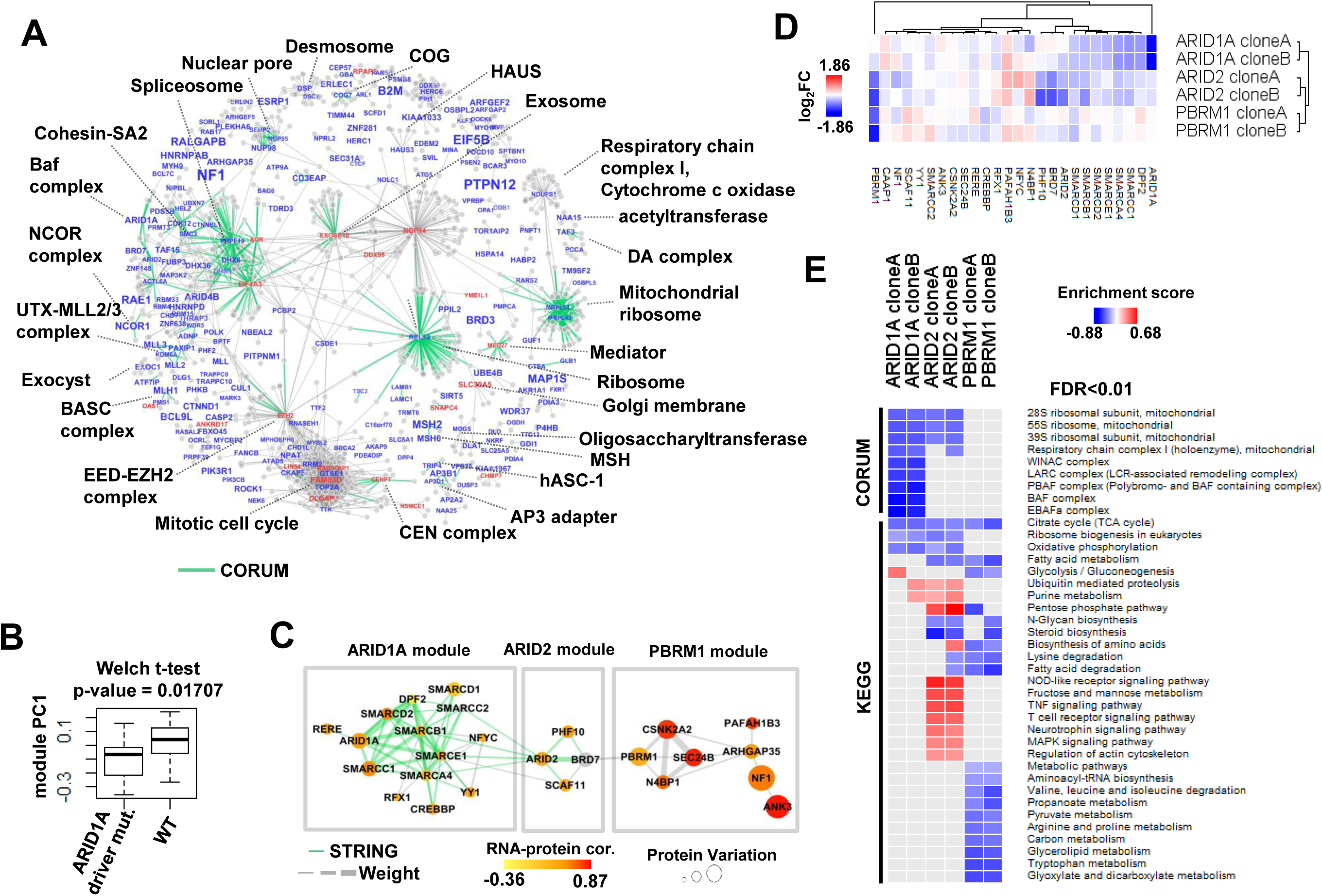
The consequences of mutations on protein complexes. A) Correlations networks filtered for known STRING interactions of proteins affected by mutations at p-value<0.05. The font size is proportional to the −log_10_(p-value) and the font colour displays the effect of the mutations on protein abundances (blue=negative, red=positive). CORUM interactions are highlighted as green thick edges and representative protein complexes are labelled. B) Boxplots illustrating the association of ARID1A driver mutations with lower levels of the ARID1A complex. C) Protein abundance correlation network of the ARID1A, ARID2 and PBRM1 modules. Green edges denote known STRING interactions and the edge thickness is increasing proportionally to the interaction weight. The node colour displays the mRNA-to-protein Pearson correlation and the size of the nodes shows the protein variation. D) Heatmap summarizing the protein abundance log_2_fold-change values in the knockout clones compared to the WT clones for the proteins in the ARID1A, ARID2 and PBRM1 modules. E) KEGG pathway and CORUM enrichment analysis for the proteomic analysis results of ARID1A, ARID2 and PBRM1 CRISPR knockouts in human iPS cells.

The above examples confirm that the loss of a subunit in a protein complex can diminish the protein abundance of its partners but not always to the same degree and with subsequent changes in the total stoichiometry. We devised linear models to detect specific mutations that cause severe deviations from strongly correlating protein profiles across the cell lines, thus significantly compromising the maintenance of stoichiometry between the interacting partners. We detected 50 such mutations (p-value<0.05); mostly frameshift or nonsense alterations but also several single amino acid substitutions (**Table S11**). Two examples are provided in **Figure S6B**. The outlier points in the correlation plots of PIK3R1-PIK3CB and SEC31A-SEC13 involved in the PI3K pathway and protein processing in endoplasmic reticulum respectively could be explained by a truncating mutation in PIK3R1 and a missense mutation in SEC31A that significantly disrupted the total abundance stoichiometry impairing their co-functionality in the associated biological processes. This analysis highlights a subset of specific mutations with the highest impact on protein abundance and reveals their cell line-specific consequences on protein interactions. Overall, our findings indicate that an additional layer of protein variation can be potentially explained by the collateral effects of mutations on tightly co-regulated partners.

### Protein quantitative trait loci analysis of colorectal cancer drivers

We performed Quantitative Trait Loci (QTL) analysis to systematically interrogate the distant effects of colorectal cancer driver genomic alterations on protein abundance (pQTL) and gene expression (eQTL). We identified 86 proteins and 196 mRNAs with at least one pQTL (**Table S12**) and eQTL respectively at 10% FDR (**Figure 5A, Figure S6C**). To assess the replication rates between independently tested QTL for each phenotype pair we also performed the mapping using 6,456 commonly quantified genes and we found that 64% of the pQTLs (N=74) validated as eQTLs and 54% of the eQTLs (N=86) validated as pQTLs (**Figure 5B**). Ranking the pQTLs (FDR<30%) by the number of associations showed that mutations in BMPR2, RNF43 and ARID1A, as well as CNAs of regions 18q22.1, 13q12.13, 16q23.1, 9p21.3, 13q33.2 and 18q21.2 loci accounted for 62% of the total variant-protein pairs (**Figure 5C**). The above-mentioned genomic events were also among the top 10 eQTL hotspots (**Figure S6D**). High frequency hotspots in chromosomes 13, 16 and 18 associated with CNAs have been previously identified in colorectal cancer tissues (Zhang et al., 2014). Enrichment analysis of the gene sets associated with each pQTLs showed overrepresentation of 12 distinct protein complexes and 36 partially redundant GO terms (Fisher’s test, Benj. Hoch. FDR<0.1). Interestingly, increased levels of the mediator complex were associated with FBXW7 mutations (**Figure S6E, first panel**), an ubiquitin ligase that targets MED13/13L for degradation (Davis et al., 2013) and TP53 mutant cell lines were associated with up-regulation of cell division related proteins (**Figure S6E, second panel**). Examination of the pQTL for other functional relationships showed that driver mutations in RNF43, an E3 ubiquitin-protein ligase that negatively regulates the Wnt signaling pathway, were positively associated with APC protein abundance (**Figure S6E, third panel**) and that BMPR2 mutations were negatively correlated with TGFBR2 protein levels (**Figure S6E, fourth panel**), both being members of the TGF-beta superfamily (Massague, 2012). Our data clearly demonstrate that a large portion of genomic variation affecting mRNA levels is not always transferred to the proteome. We see that distant protein changes attributed to variation in cancer driver genes can be regulated directly at the protein level and are not conspicuous at the mRNA level, with indication of further causal effects including enzyme substrate relationships.

**Figure 5.**
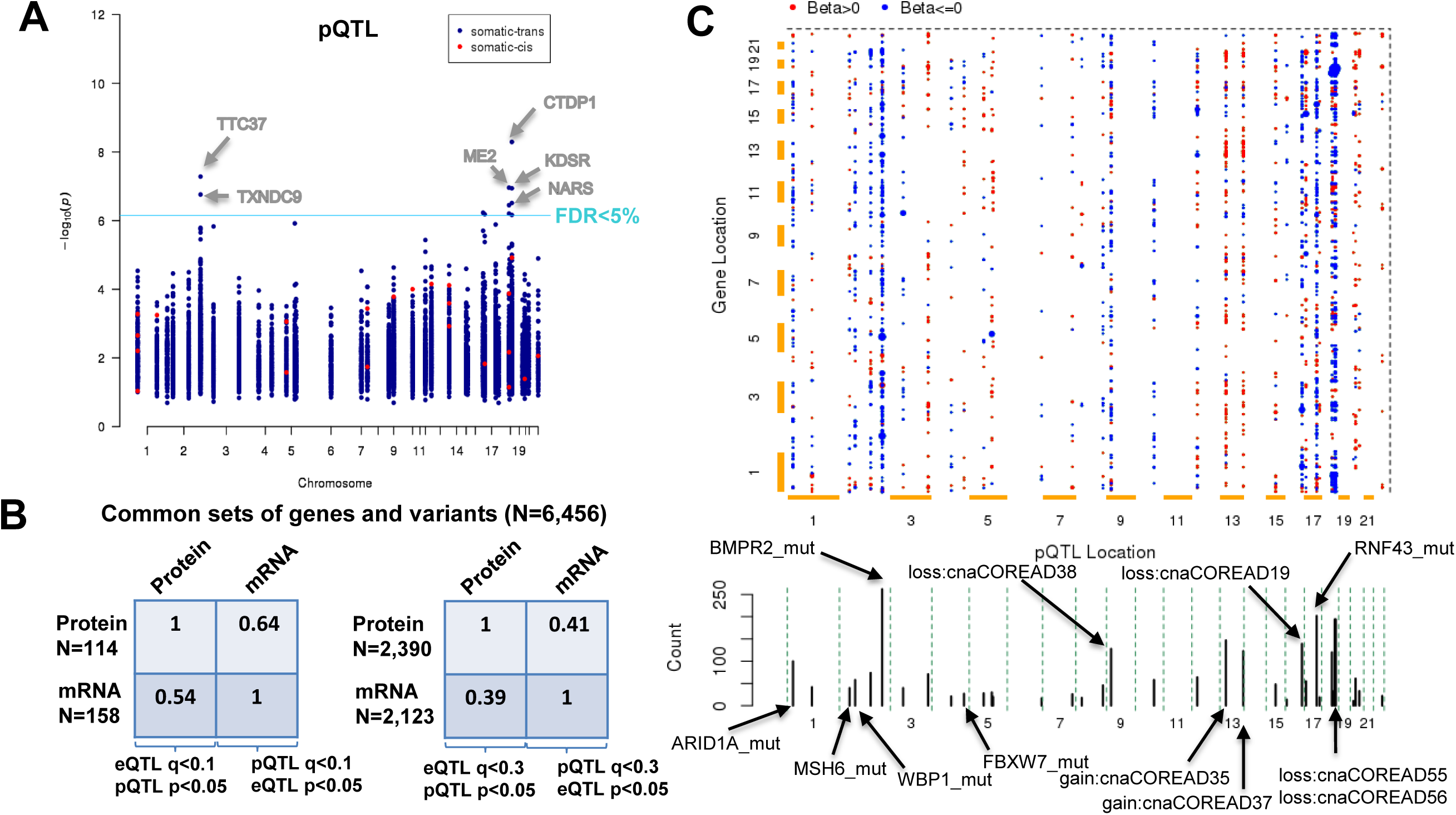
Proteome-wide quantitative trait loci (pQTL) analysis of cancer driver genomic alterations. A) Identification of cis and trans pQTLs in colorectal cancer cell lines considering colorectal cancer driver variants. The p-value and genomic coordinates for the most confident non-redundant protein-variant association tests are depicted in the Manhattan plot. B) Replication rates between independently tested QTL for each phenotype pair using common sets of genes and variants (N=6,456 genes). C) Representation of pQTLs as 2D plot of variants (y-axis) and associated genes (x-axis). Associations with q<0.3 are shown as dots coloured by the beta value (red: positive association, blue: negative association) while the size is increasing with the confidence of the association. Cumulative plot of the number of associations per variant is shown below the 2D matrix.

### Protein complexes associated with microsatellite instability

Loss of DNA mismatch repair activity is responsible for the microsatellite instability (MSI) observed in 15% of all colorectal cancers. MSI tumours are associated with better prognosis and differential response to chemotherapy (Boland and Goel, 2010). An improved understanding of the effect of MSI on cellular processes thus has the potential to explain some of these clinical features. We detected 10 differentially regulated modules between the MSI-high and MSI-low cell lines (Welch’s t-test, permutation based FDR<0.05). This encompasses 172 proteins (**Figure 6A**) that include a subset of 33 genes previously attributed to MSI events in colorectal tumors (Kim et al., 2013) such as MSH6, MSH3, PMS2, BAX and RAD50. The STRING interactions between the MSI associated proteins are depicted in **Figure 6B**, substantiating the functional relationships between this set of proteins. We additionally identified epigenetic dysregulation characterized by reduced levels of histone methyltransferase (KMT2D, PAXIP1, NCOA6, SETD1B, KMT2C, KMT2B and KDM6A). We also detect histone deacetylation (HDAC3 and NCOR1) protein complexes associated with these events. This network indicates the suppression of two members of the INO80 chromatin remodelling complex (INO80D and ACTL6A) in MSI-high cells and can explain the down-regulation of the Arp2/3 protein complex by protein interactions with ACTL6A. Other distinct complexes we have identified negatively affected by MSI were the exocyst complex, which is implicated in targeting secretory vesicles to specific docking sites on the plasma membrane (Heider and Munson, 2012) and the SKI complex (SKIV2L, TTC37 and WDR61), which is involved in exosome-mediated RNA decay (Wang et al., 2005). Overall, this alludes to multiple epigenetic mechanisms playing a role in the MSI pathogenesis, and also suggests a new role for exocyst in this phenotype. The MSI up-regulated modules showed over-representation of proteins from the loci 8p21 and 18q21 including SMAD4. Although high SMAD4 levels have been previously associated with MSI and better prognosis in colon cancer (Isaksson-Mettavainio et al., 2012), our data suggest that the observed differences stem from a mutually exclusive SMAD4 copy number alteration (loss) with MSI.

**Figure 6.**
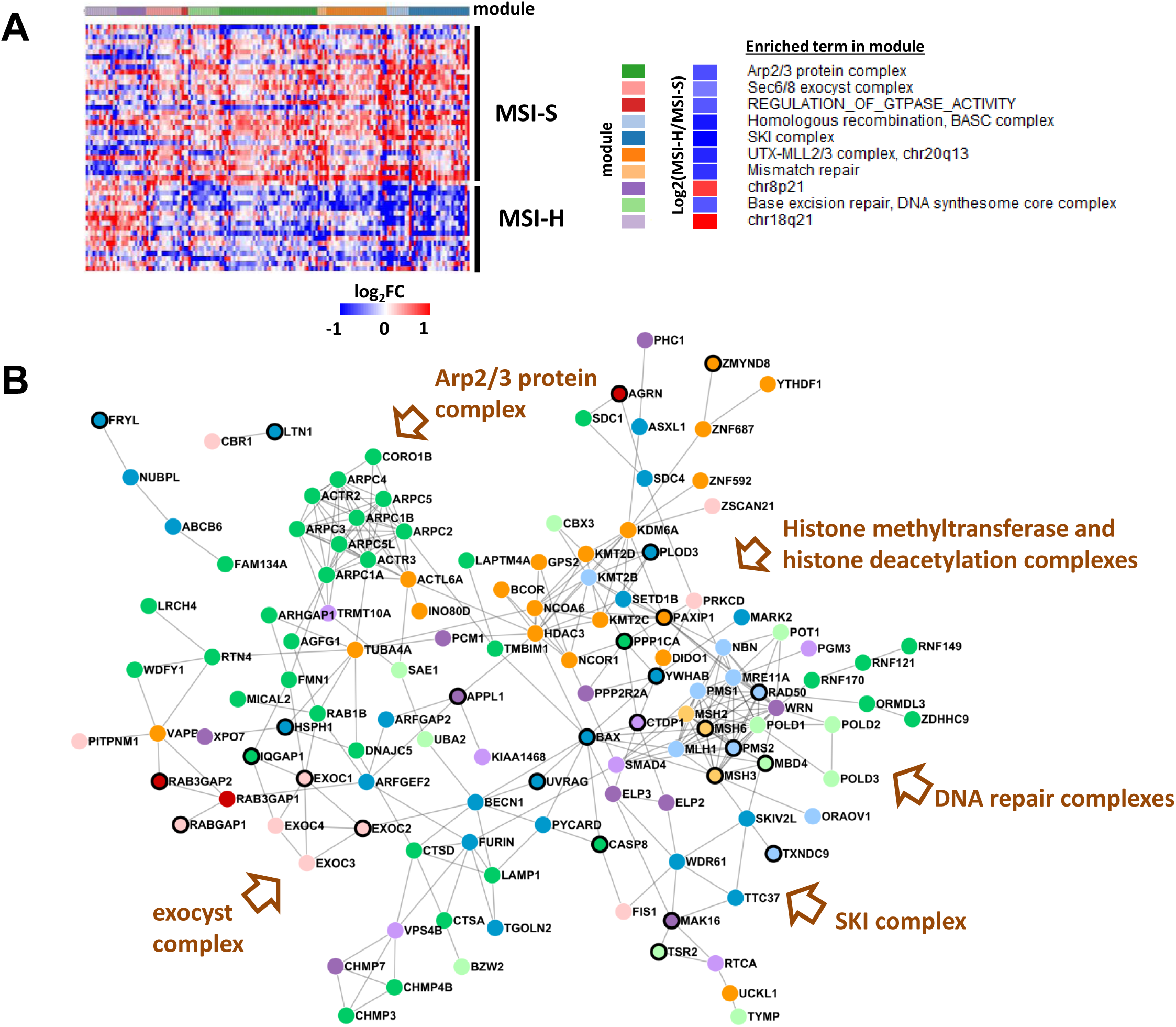
Protein complexes associated with MSI. A) Heatmap of MSI-high associated proteins (Welch t-test, permutation based FDR<0.05). Columns represent proteins sorted horizontally based on color-coded modules and rows correspond to cell lines. The modules are labelled by significantly enriched terms on the right panel. B) STRING network of interconnected MSI-high associated proteins. The nodes are color-coded by module and distinct complexes are highlighted. Nodes with black outline have been previously found with MSI events in colorectal cancer by Kim et al.

### Proteomic subtypes of colorectal cancer cell lines

To explore whether our deep proteomes recapitulate tissue level subtypes of colorectal cancer and to provide insight into the cellular and molecular heterogeneity of the colorectal cancer cell lines, we performed unsupervised clustering based on the quantitative profiles of 7,330 proteins without missing values by class discovery using the ConsensusClusterPlus method (Wilkerson and Hayes, 2010). Optimal separation by k-means clustering was reached using 7 colorectal proteomic subtypes (CPS) (**Figure S7A and Figure 7A**).

**Figure 7.**
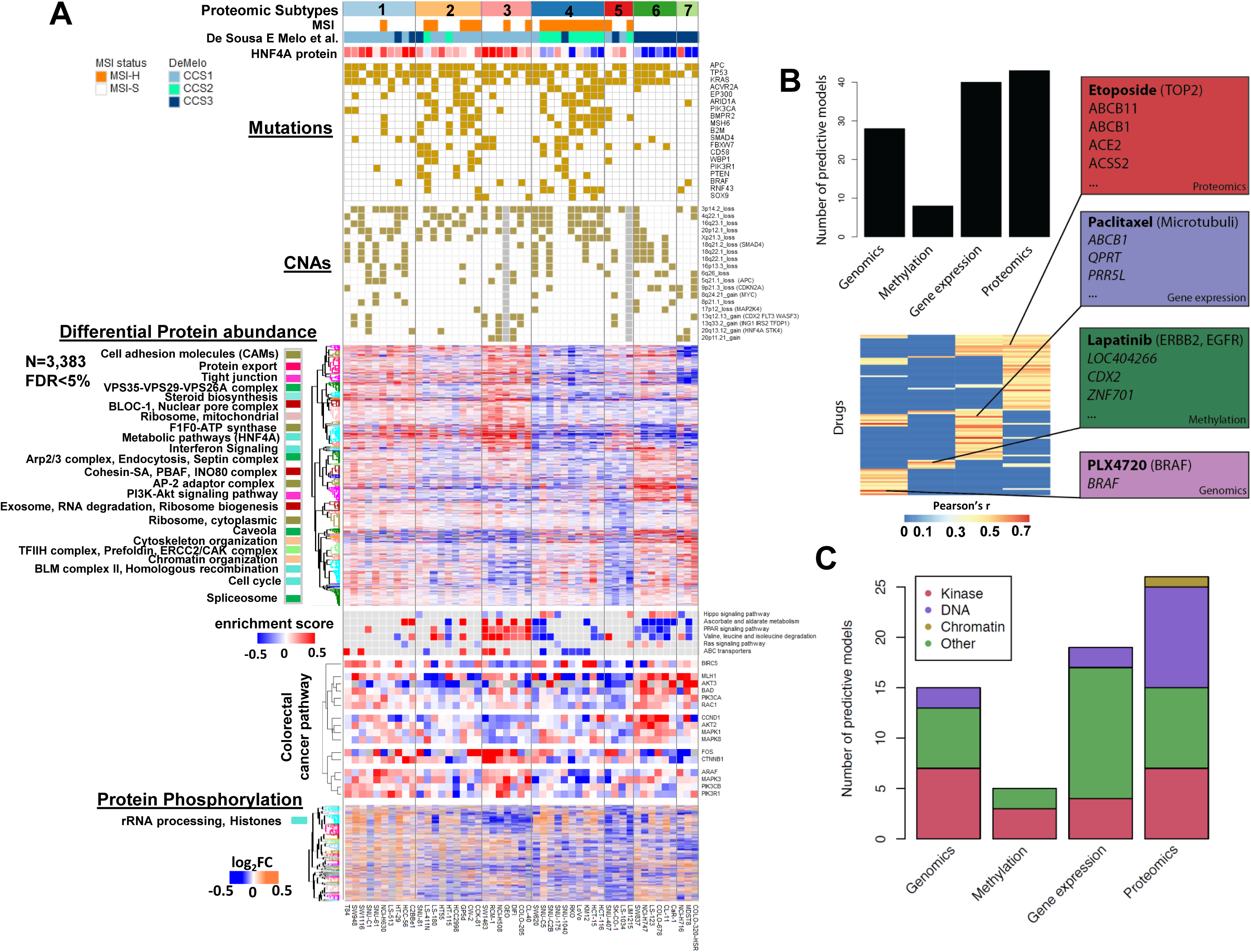
Proteomics subtypes of colorectal cancer cell lines and drug associations. A) Cell lines are represented as columns, horizontally ordered by seven color-coded proteomics consensus clusters and aligned with microsatellite instability (MSI), published colorectal cancer subtypes by DeMelo classification, HNF4A protein abundance, cancer driver genomic alterations, differentially regulated proteins, selected enriched KEGG pathways, differentially regulated colorectal cancer proteins and differentially regulated phosphopeptides. The heatmap of the differentially regulated proteome was divided into 50 color-coded clusters. Enriched terms for each cluster are shown on the left. B) The number of drugs for which predictive models (i.e. models where the Pearson correlation between predicted and observed IC50s exceeds r > 0.4) could be fitted is stratified per data type (top panel). Predictive models for more drugs are found by the use of proteomics data. A heatmap indicating for each drug and each data type whether a predictive model could be fitted (bottom panel). Drugs for which no predictive model could be fitted using any data type were omitted. Most drugs were specifically predicted by one data type. Examples of predictive models for each of the four data types are highlighted. C) The number of drugs where response was specifically predicted by one molecular data type stratified by each of the four molecular data types and by four classes defined based on drug target and biological activity.

Our proteomic clusters overlapped very well with previously published tissue subtypes (annotations from Medico et al., **Figure S7B**) (Medico et al., 2015), especially with the classification described by De Sousa E Melo et al. (De Sousa E Melo et al., 2013). Previous classifiers have commonly subdivided samples along the lines of ‘Epithelial’ (lower crypt and crypt top), ‘MSI-H’ and ‘Stem-like’, with varying descriptions (Guinney et al., 2015). In contrast, our high depth proteomic dataset not only captured the commonly identified classification features, but provides increased resolution to further subdivide these groups. The identification of unique proteomic features pointing to key cellular functions, gives insight into the molecular basis of these subtypes, and provides clarity as to the differences between them (**Figure 7A**). A detailed description of the unique proteomic features of our COREAD classification is provided in **Table S13**.

Cell lines with a canonical epithelial phenotype (previously classified as CCS1 by De Sousa E Melo et al., 2013) clustered together, but are now divided into 3 subtypes (CPS1, CPS2, CPS3). These subtypes all displayed high expression of HNF4A, indicating a more differentiated state. While subtypes CPS1 and CPS3 contain Transit Amplifying cell phenotypes (Sadanandam et al., 2013), CPS2 is largely characterised by a Goblet cell signature (**Figure S7B**). CPS2 is also enriched in lines that are hypermutated, and while some are MSI-H, the MSI-negative/hypermutated lines (HT115, HCC2998, HT55) (Medico et al., 2015) all cluster in this group (**Figure S7B**). Transit Amplifying subtype CPS3 can be distinguished from CPS1 by lower expression of cell cycle proteins (eg. CCND1), and low histone phosphorylation (possibly mediated by VRK3), as well as higher activation of PPAR signalling and amino-acid metabolism pathways. CPS3 also contains lines (DIFI, NCI-H508) that are most sensitive to the anti-EGFR antibody Cetuximab (Medico et al., 2015). Further, this group correlates with a crypt top description ‘Subtype A’ (Budinska et al., 2013) while subtypes CPS1 and CPS2 are associated with the lower crypt ‘Subtype B’ (Budinska et al., 2013), (**Figure S7B**).

The CPS4 subtype is the canonical MSI-H cluster, with a strong correlation with the CCS2 cluster identified by De Sousa E Melo et al. These lines have also been commonly associated with a less differentiated state by other classifiers, and this is reinforced by our dataset; subtype CPS4 has low levels of the HNF4A-CDX2 module, rendering this group clearly distinguishable from CPS2 (**Figure 7A**). The separation into two distinct MSI-H/Hypermutated classifications was also observed by Guinney et al., (2015), and may have implications for patient therapy and prognosis. Significantly, CPS4 displays low expression of ABC transporters, which may contribute to the better response rates seen in MSI-H patients (Popat et al., 2005).

The origin of CPS5 is less well defined, as it expresses intermediate levels of HNF4A. However, it is characterized by moderate down-regulation of cell cycle, ribosome and spliceosome modules, and displays low levels of MLH1 and AKT3 proteins (**Figure 7A**). Interestingly, the phosphorylation landscape of CPS5 exhibits a global low phosphorylation, particularly in microtubule cytoskeleton and adherens junction proteins.

Lastly, we capture the commonly observed colorectal ‘Stem-like’ subgroup, which is represented in subtypes CPS6 and CPS7 (**Figure 7A, S7B**). Both subtypes exhibit stem-like expression profiles, with very low levels of HNF4A and CDX1 transcription factors (Chan et al., 2009; Garrison et al., 2006; Jones et al., 2015). Cells in both subtypes commonly exhibit loss of 9p21.3 including *CDKN2A* and *CDKN2B,* while this is rarely seen in other subtypes. Interestingly, while CPS6 displays activation of the Hippo signalling pathway and loss of 18q21.2 *(SMAD4),* CPS7 has a mesenchymal profile, with low expression of CDH1 and AXIN2, and high Vimentin. The overall strong suppression of cell adhesion and tight junction components may be influenced by low expression of KLF5 (Zhang et al., 2013).

### Pharmacoproteomic models are strong predictors of response to DNA damaging agents

Although a number of recent studies have investigated the power of different combinations of molecular data to predict drug response in colorectal cancer cell lines, these have been limited to using genomic (mutations and copy number), transcriptomic and methylation datasets (Iorio et al., 2016). We have shown above that the DNA and gene expression variations are not perfectly mirrored in the protein measurements. As such one might expect to gain predictive power for some phenotypic associations when also using the protein abundance changes. To date there has not been a comprehensive analysis of the effect on the predictive power from the addition of proteomics datasets in colorectal cancer. All of the colorectal cell lines included in this study have been extensively characterised by sensitivity data (IC50 values) for 265 compounds (Iorio et al., 2016). These include clinical drugs (n = 48), drugs currently in clinical development (n = 76), and experimental compounds (n = 141).

We built Elastic Net models that use as input features genomic (mutations and copy number gains/losses), methylation (CpG islands in gene promoters), gene expression and proteomic datasets. We were able to generate predictive models where the Pearson correlation between predicted and observed IC50 >0.4 in 91 of the 265 compounds (**Table S14**). Importantly, using the proteomics data enabled the construction of more predictive models than with any other feature type (**Figure 7B, top panel**). Examples of the proteomics predictive models for etoposide are shown in **Figure S7C**. Response to most drugs was often specifically predicted by one data type, with very little overlap (**Figure 7B bottom panel**). Interestingly, when response to a drug was predicted by both gene expression and proteomics, the protein-RNA correlation for genes associated with response tended to be higher (Mann-Whitney U test, p-value: 0.006) (**Figure S7D**).

Within the proteomics-based signatures found to be predictive for drug response, we frequently observed the drug efflux transporters ABCB1 and ABCB11 (8 and 7 out of 43 respectively, 9 unique) (**Table S14**). In all models containing these proteins, elevated expression of the drug transporter was associated with drug resistance. Interestingly, ABCB1 and ABCB11 are very tightly co-regulated (Pearson’s r=0.94, FDR= 5.99E-21), suggesting a novel interaction. Notably, protein measurements of these transporters correlated more strongly with response to these drugs than the respective mRNA measurements (mean Pearson’s r=0.68 and r=0.43 respectively, Wilcoxon test p-value=0.0005). This suggests that the protein expression levels of drug efflux pumps play a key role in determining drug response, and while predictive genomic biomarkers may still be discovered, the importance of proteomic associations with response should not be under-estimated.

To detect whether any specific classes of drug might be best predicted by each of the 4 molecular features, we initially classified each of the 91 agents into 21 classes depending on target class and biological activity (**Figure S7E**), and subsequently further reduced the dimensionality of the data by then classifying into 4 groups (termed ‘kinase’, ‘DNA’, ‘chromatin’ and ‘other’). There was a significant enrichment for proteomics within the predictive models for the ‘DNA’ group of agents, which includes many chemotherapy agents and mitotic poisons (Mann-Whitney U test, nominal p-value: 0.01, FDR-corrected p-value: 0.0486) (**Figure 7C**). In contrast, 47% of the models specifically predicted using genomics features were for kinase inhibitors. This suggests that while the response to targeted kinase inhibitors can be modulated by point mutation (eg. BRAF mutations predict response to BRAF inhibitors), the response to broader DNA damaging agents is dependent on the expression levels of key proteomic subsets. Interestingly, NBEAL1 and PARD3B proteins that were found to be down-regulated by mutations were also among the top 10 proteomics predictive models, were absent from the gene expression models, and 6 out of the 11 associated drugs were from the "DNA” category. Although the mechanism by which these proteins may affect drug sensitivity are unclear, it is known that PARD3B is involved in asymmetrical cell division and cell polarization processes (Williams et al., 2014) and that both genes are located in chr2q33 and are associated with Amyotrophic Lateral Sclerosis 2 which suggests functional similarities. These examples also highlight the value of proteomics in better understanding the consequences of non-driver genomic alterations in drug sensitivity through the proteome.

## Discussion

Our analysis of colorectal cancer cells using in-depth proteomics has yielded several significant insights into both fundamental molecular cell biology, and the molecular heterogeneity of colorectal cancer subtypes. Beyond static measurements of protein abundances, the quality of our dataset enabled the construction of a reference proteomic co-variation map with topological features reflecting the dynamic interplay between protein complexes and biological processes in colorectal cancer cells. Notably, identification of protein complexes and network topologies in such a global scale would require the analysis of hundreds of protein pull-downs and thousands hours of analysis (Hein et al., 2015) thus our approach can serve as a time effective screening tool for the study of protein networks. Another novel aspect that emerged from our analysis is the maintenance of co-regulation at the level of net protein phosphorylation. This seems to be more pronounced in signalling pathways where the protein abundances are insufficient to indicate functional associations. Analogous study of co-regulation between different types of protein modifications could also enable the identification of modification cross-talk (Beltrao et al., 2013).

We show that the subunits of protein complexes tend to tightly maintain their total abundance stoichiometry post-transcriptionally which forms the basis for the better understanding of the higher order organization of the proteome. The primary level of co-regulation between proteins allows for prediction of human gene functions and the secondary assembly of the co-variome reveals the interdependencies of protein complexes and biological processes and uncovers possible pathway interplays. Importantly, our data can be used in combination with genetic interaction screens (Costanzo et al., 2016) to explore whether gene essentiality meets the protein co-regulation principles. Our catalogue of 210,000 weighted interactions can help the selection of protein hubs representing the best predictors of interactomes in pull-down assays. Moreover, the identification of proteins with outlier profiles from the conserved profile of their known interactors, within a given complex, can point to their pleiotropic roles in the associated processes.

The simplification of the complex proteomic landscapes enables a more direct alignment of genomic features with cellular functions and delineates how genomic variation is received by protein networks and how this is disseminated throughout the proteome. This framework also proved very efficient to identify upstream regulatory events that link transcription factors to their transcriptional targets at the protein level and explained the components of the co-variome not strictly shaped by physical protein interactions. To a smaller degree the module-based analysis was predictive of co-occurring genomic variants exposing paradigms of simple cause-and-effect proteogenomic features of the cell lines.

We show that mutations largely affect protein abundances directly at the protein level with a higher pressure on the chromatin modification protein class. Targeted depletion of key chromatin modifiers by CRISPR/cas9 followed by proteomic analysis confirmed that the effects of genomic variation on distant gene products, physically related with the directly affected proteins, can be explained by the mechanisms that define protein co-variation. The latter is supported by the observation that the severity of the distant effects is well predicted by the co-variation consistency and the topological features of the correlation networks. Additionally, this analysis indicated that directionality can be another characteristic of such interactions.

We provide evidence that colorectal cancer subtypes derived from tissue level gene expression datasets are largely reproduced at the proteome level which further resolves the main subtypes into groups that reflect a possible cell type of origin and the underlying differences in genomic alterations. This robust functional characterization of the COREAD cell lines can be a useful resource to guide cell line selection in targeted cellular and biochemical experimental designs where cell line specific biological features can bias the results. Importantly, proteomics analysis highlighted that protein variation better predicts responses to drugs that interfere with cell cycle and DNA replication and that the expression of key protein components such as ABC transporters is critical to predicting drug response in colorectal cancer. While further work is required to establish these as validated biomarkers of patient response in clinical trials, numerous studies have noted the role of these channels in aiding drug efflux (Chen et al., 2016). Overall, our Elastic Net models suggest that expression of a single protein alone may not be sufficient to predict drug resistance, and consideration of a panel of markers may be required. This study demonstrates that proteomics is the technology of choice for functional systems biology and provides a valuable resource for the study of regulatory variation in cancer cells.

## Author Contributions

Conceptualization, J.S.C. & U.M.; Methodology, T.I.R., S.W.; Mass Spectrometry T.I.R.; Proteomics Data Analysis, T.I.R., E.G., M.S., S.W., J.C.W., P.B., J.S-R.; QTL Analysis, F.Z.G., E.G., A.B., J.S-R.; O.S.; Drug Data Analysis, N.A., M.M., M.S., M.Y., J.S-R.; S.W., T.I.R., L.W., U.M.; CRISPR Lines, C.A., D.J.A.; Writing–Original Draft, T.I.R., S.W., L.W., U.M. J.S.C.; Writing–Review & Editing, All.

## Acknowledgments

This work was funded by the Wellcome Trust (086375 and 102696). SPW is funded by the ERC Synergy Project CombatCancer. We would like to thank members of the Cancer Genome Project for helpful discussions and Sarah A. Teichmann for discussions and general suggestions about the manuscript.

## Supplementary figures legends

**Figure S1.**
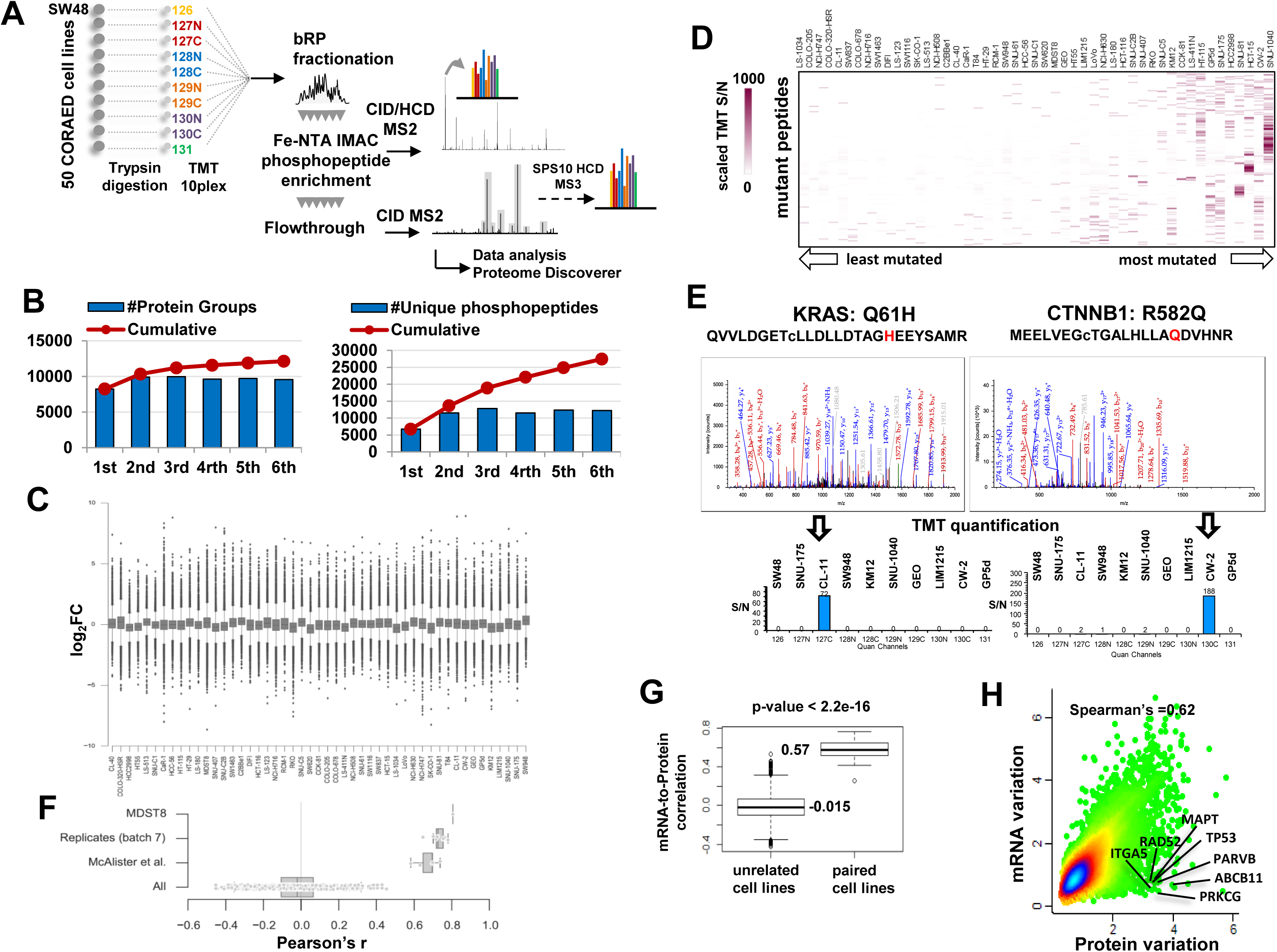
Proteome and phosphoproteome coverage. A) Workflow for quantitative global proteome and phosphoproteome analysis. 50 colorectal cancer cell lines (COREAD) were analysed using TMT-10plex in seven multiplex sets. The SW48 cell line was used as the reference sample in each set. Biological replicates of MDST8 cell line were included in two different sets and the 7th set corresponds to a biological replicate of the 6th set. These were used to evaluate the normalization and the batch effect correction methods. B) Number of protein groups (left panel) and unique phosphopeptides (right panel) identified per multiplex set are depicted as blue bars and cumulative number of identifications shown as red lines. C) Box plots of normalized log_2_Ratio values per cell line. D) Heatmap of the TMT scaled S/N values of the identified mutant peptides shown as rows. The columns represent the COREAD cell lines sorted from the least mutated (left) to the most mutated (right) cell line. E) Two example identification spectra of mutant peptides from KRAS and CTNNB1 along with the TMT quantification profiles. F) Box plots of Pearson correlation coefficients between un-related samples (All), paired samples from the 7th replicate batch, the MDST8 replicate samples and inter-laboratory replicates for six cell lines from McAlister et al. G) Box plots of the mRNA to protein Pearson correlation between unrelated and paired cell lines. H) Scatter plot of protein variation versus mRNA variation expressed as median-normalized standard deviation (SD) across the cell lines.

**Figure S2.**
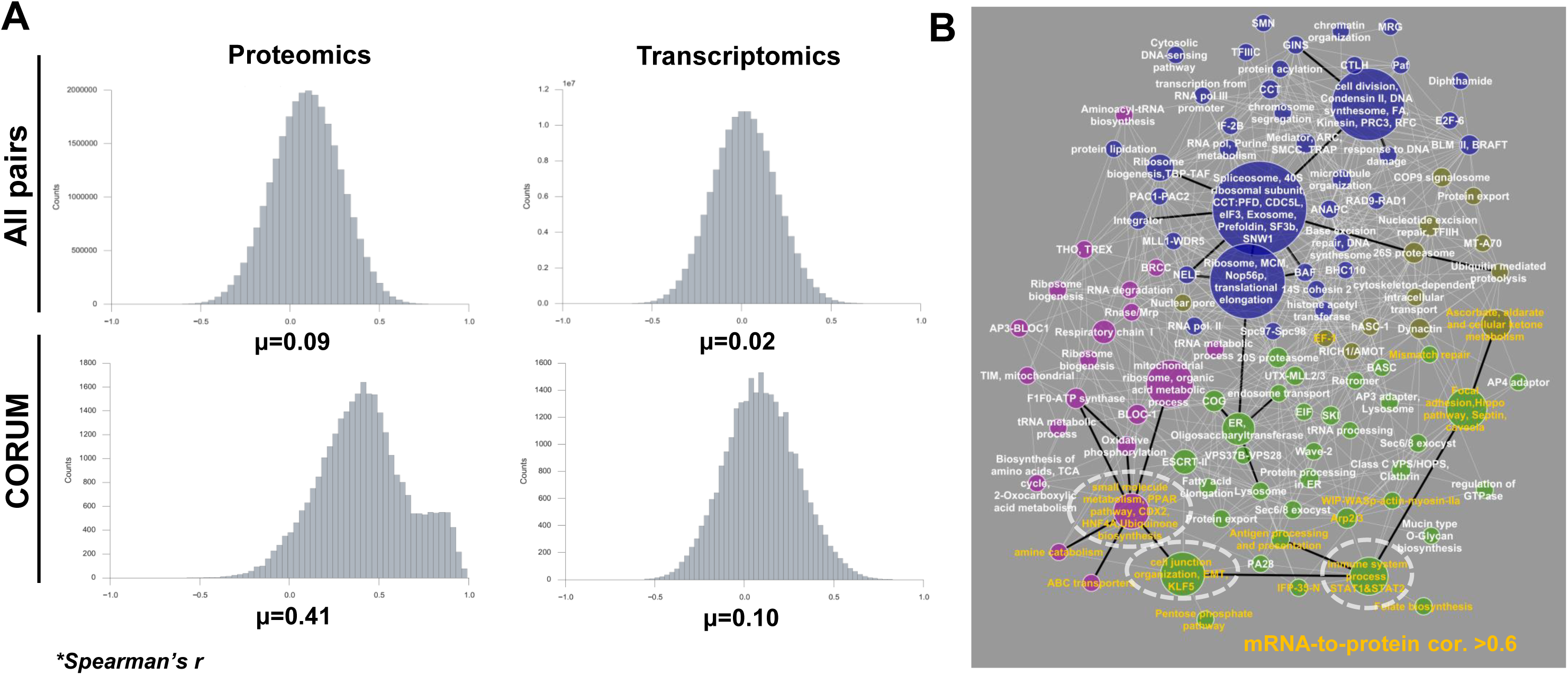
Global distributions of gene-to-gene correlations. A) Distributions of Spearman’s correlation coefficients between protein-protein pairs (left panel) and mRNA-mRNA pairs (right panel) for all pairs and for pairs with known relationships in the CORUM database. B) Correlation network of the WGCNA modules using the eigengene profiles (Pearson>0.46, Benj. Hoch. FDR<0.01). Nodes represent WGCNA modules labelled with enriched terms (GO-Slim, KEGG, CORUM, GSEA, ChEA, ENCODE and Pfam) and are color-coded by ReactomeFI clusters. The size of the nodes is proportional to the number of proteins in the module. Transcriptionally controlled processes are highlighted with orange font and the processes with enriched transcription factors are outlined. Black thick edges highlight examples of associations between biological processes or protein complexes.

**Figure S3.**
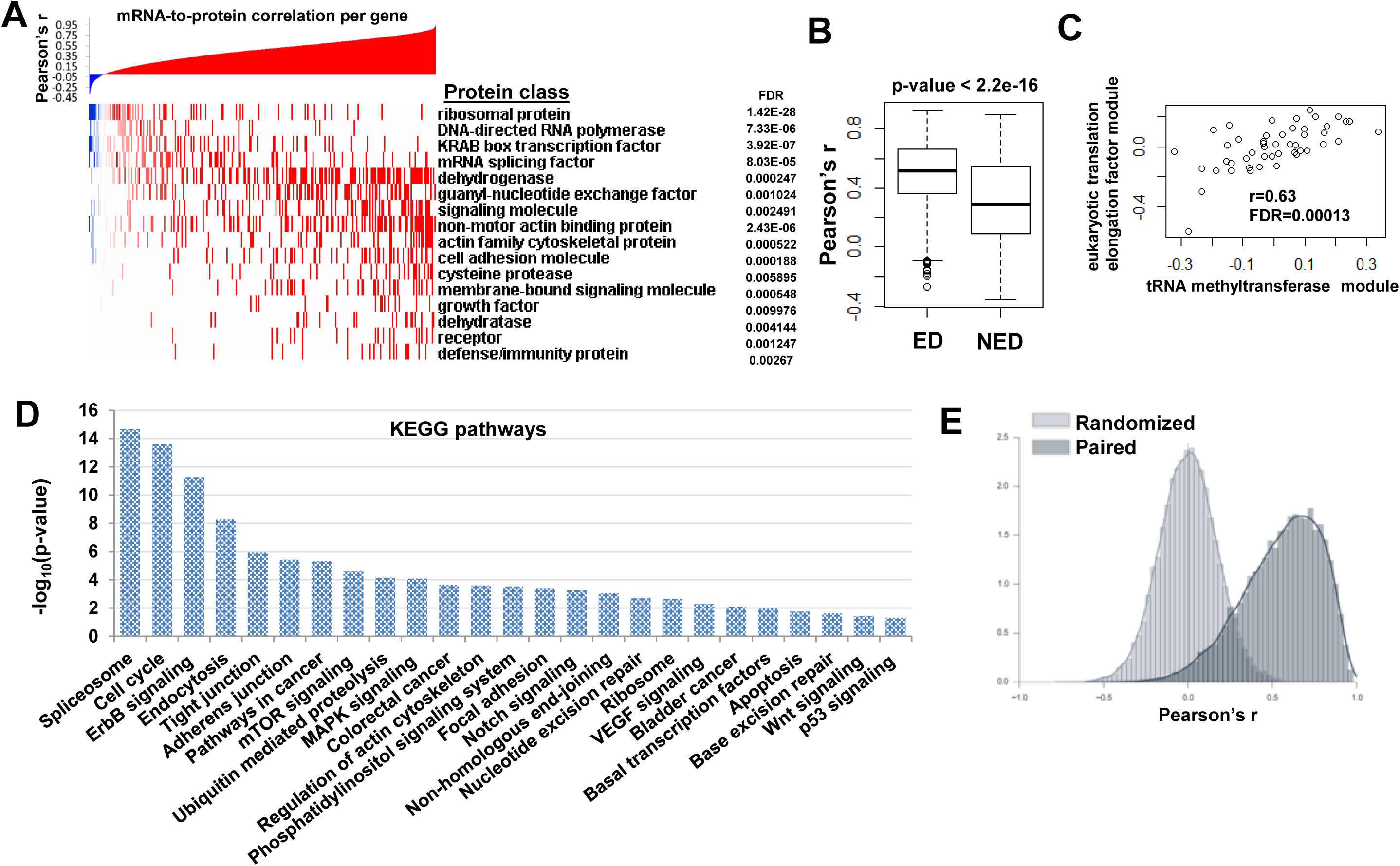
Transcriptome-to-proteome correlation per protein class, protein modules correlation plot and phosphorylated pathways. A) Gene-level mRNA-to-protein Pearson correlations ranked by lowest to highest value. PANTHER protein classes with negative or positive enrichment relatively to the mean of all mRNA-to-protein correlations are displayed. B) Box plots illustrating the mRNA-to-protein correlation for proteins characterized as exponentially degraded (ED) and non-exponentially degraded (NED) by McShane et al., 2016. C) Scatter plot illustrating the correlation between tRNA methyltransferases and eukaryotic translation elongation factors. D) Enriched KEGG pathways by DAVID analysis of all quantified phosphoproteins. E) The distributions of Pearson coefficients for randomized and matched pairs of phosphopeptide abundances versus protein abundances.

**Figure S4.**
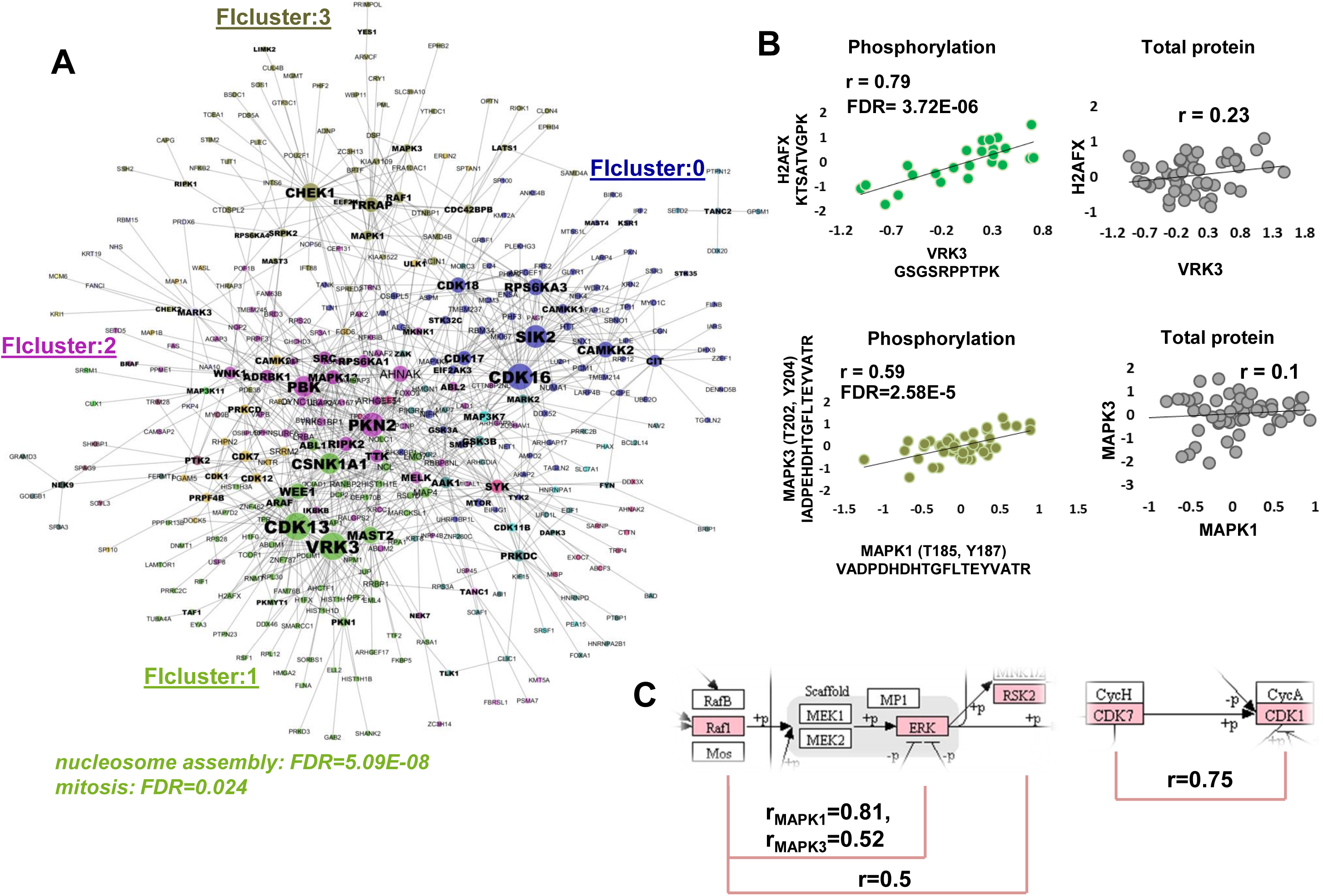
*De novo* prediction of phosphorylation networks. A) Correlation network of 213 variable phosphopeptides (with protein abundance regressed out) belonging to 144 non-redundant kinases and the 787 most variable phosphopeptides of all other types of proteins (significant Pearson correlations displayed in the network were filtered for Benjamini-Hochberg adjusted p-value<0.05). The nodes are enlarged proportionally to the number of direct edges and are color-coded based on ReactomeFI clustering. Protein kinases are highlighted with bold font. B) Scatter plots of two significantly correlating phosphoprotein pairs (VRK3-H2AFX and MAPK1-MAPK3) for which the respective protein levels displayed insignificant correlation. C) Snapshots of the MAPK and cell cycle KEGG pathways highlighting (pink) significantly correlating phosphorylations. The Pearson correlation for each association is shown below the pathways.

**Figure S5.**
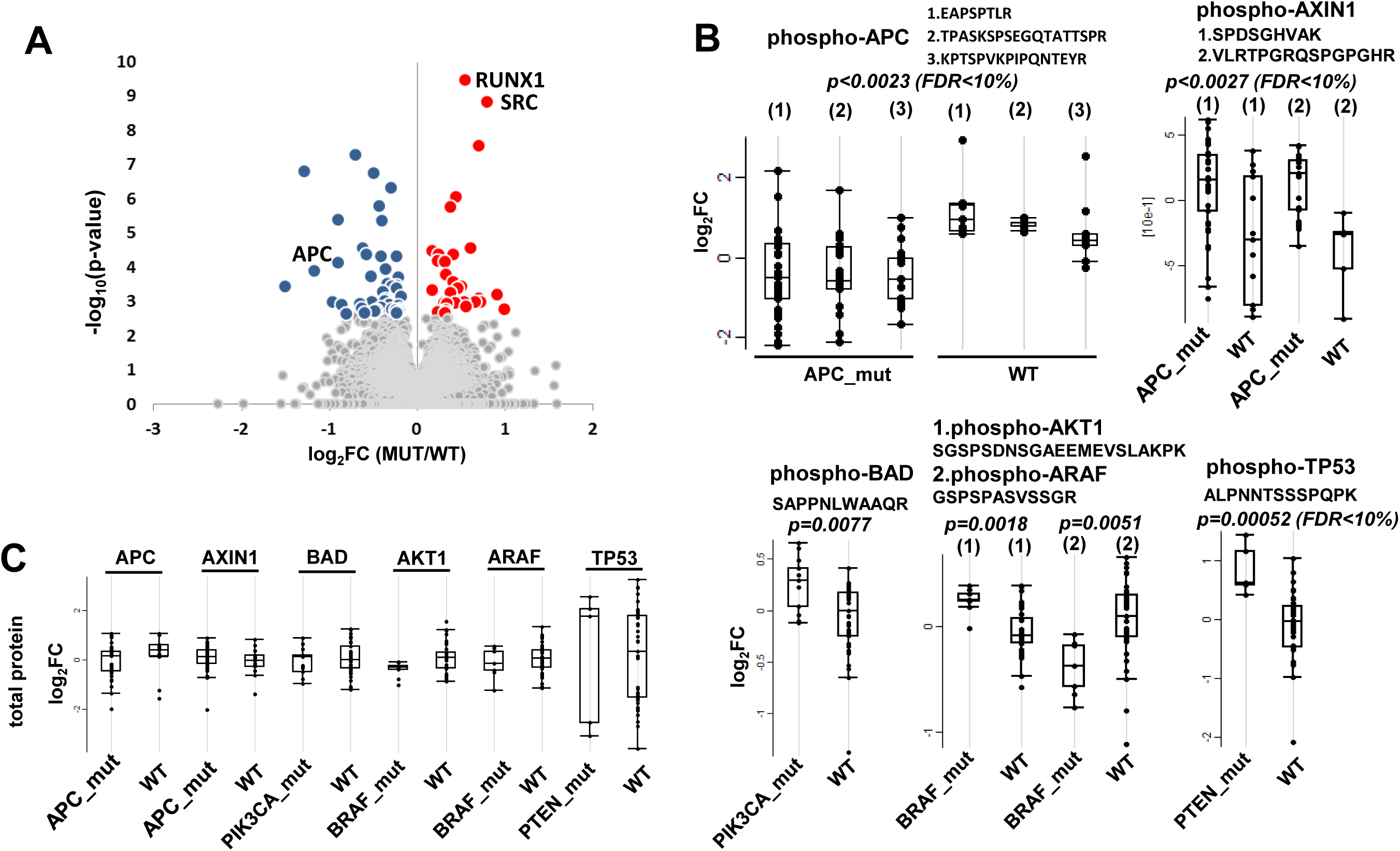
The effects of mutations on protein phosphorylation. A) Volcano plot summarizing the direct effects of mutations on protein phosphorylation. B) Box plots illustrating the differential phosphorylation between mutated and wild-type cells considering colorectal cancer driver mutations. C) The respective un-differentiated protein abundances.

**Figure S6.**
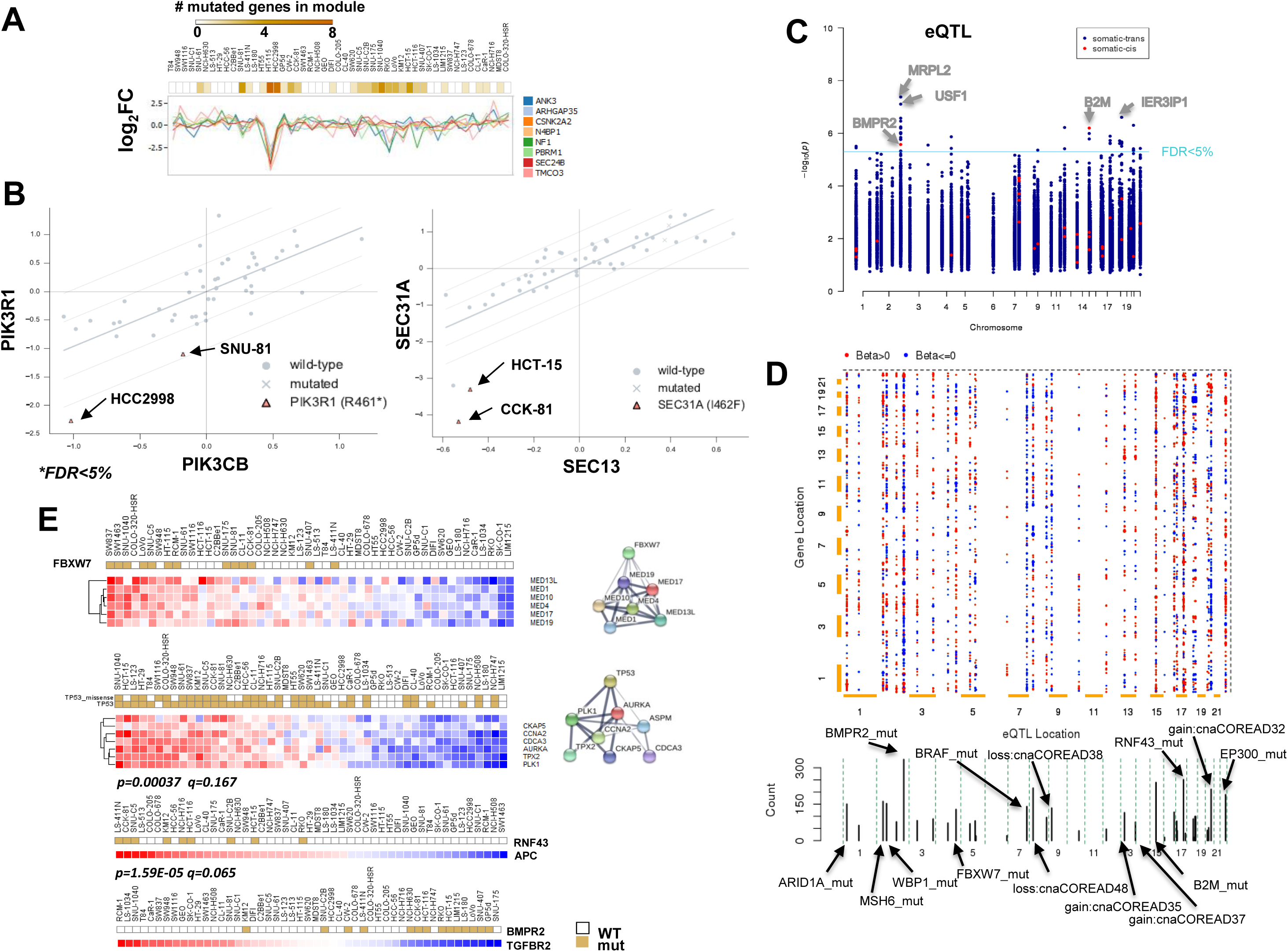
Identification of mutations that cause loss of correlation stoichiometry and expression quantitative trait loci analysis of cancer driver genomic alterations. A) Line plot displaying the abundance profile of the proteins in the PBRM1 module. The top bar indicates the total number of mutated genes within the module across the cell lines. B) Scatter plots of protein pairs in which specific mutation cause severe divergence from highly correlating profiles. C) Identification of *cis* and *trans* eQTLs in colorectal cancer cell lines considering cancer driver variants. The p-value and genomic coordinates for the most confident non-redundant mRNA-variant association tests are depicted in the Manhattan plot. D) Representation of eQTLs as 2D plot of variants (y-axis) and associated genes (x-axis). Associations with q<0.3 are shown as dots coloured by the beta value (red: positive association, blue: negative association) while the size is increasing with the confidence of the association. Cumulative of the number of associations per variant is plotted below the 2D matrix. E) Selected examples of protein networks and individual proteins with pQTLs functionally associated with the cancer variants.

**Figure S7.**
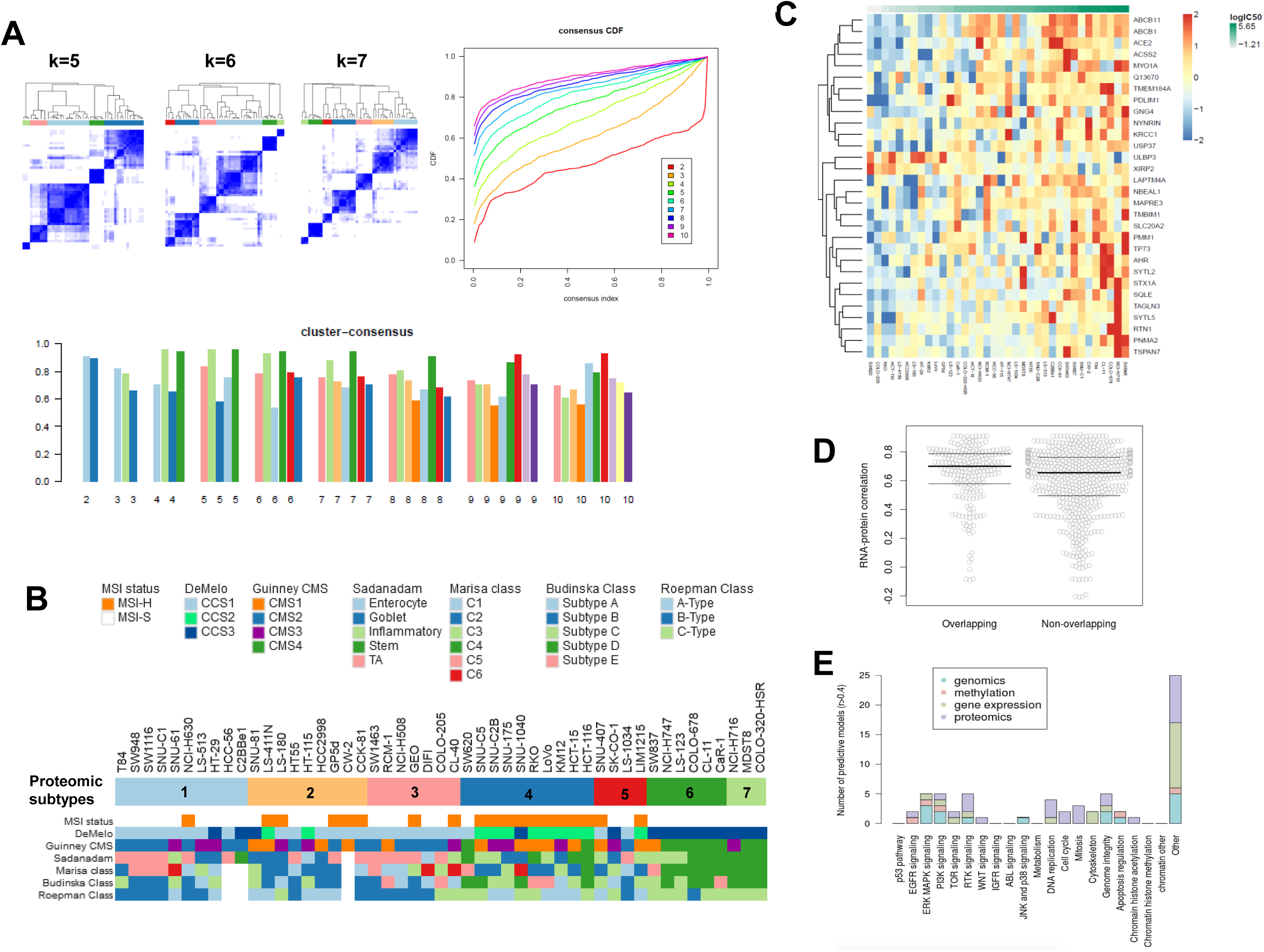
Consensus clustering of colorectal cancer cell lines and drug response models for each drug target class and for each data type. A) Proteome clusters were derived based on consensus clustering. Optimal classification of the cell lines based on the proteome quantified across all cell lines (N=7,330) was derived by the application of the ConsensusClusterPlus R package using 1,000 resampling repetitions in the range of 2 to 10 clusters. The consensus matrices for target values *k*=5,6 and 7 are visualized (top left panel) along with the empirical cumulative distribution function (CDF) plot which indicates the *k* at which the distribution reaches an approximate maximum (top right panel). Cluster-consensus plot displaying the mean of all pairwise consensus values between a cluster’s members at each *k*. Balanced mean consensus values are obtained at *k*=7. B) Overlap of the proteomics subtypes with tissue level classifications. C) Heatmap showing the proteomic signature associated with response to Etoposide. D) Gene-level mRNA-to-protein Pearson correlation for genes associated with drug response, for drugs that could be predicted by both gene expression and proteomics data (overlapping) or for drugs that could only be predicted gene expression or proteomics (non-overlapping). E) The number of drugs where response was specifically predicted by one molecular data type, stratified by each of the four molecular data types and by the 21 drug classes defined by Iorio et al. (2016).

